# BRCA1 Mediated Homologous Recombination and S Phase DNA Repair Pathways Restrict LINE-1 Retrotransposition in Human Cells

**DOI:** 10.1101/701458

**Authors:** Xiaoji Sun, Paolo Mita, David J. Kahler, Donghui Li, Aleksandra Wudzinska, Chi Yun, Joel S. Bader, David Fenyö, Jef D. Boeke

## Abstract

Long interspersed element-1 (LINE-1 or L1) is the only autonomous retrotransposon active in human cells. L1s DNA makes about 17% of the human genome and retrotransposition of a few active L1 copies has been detected in various tumors, underscoring the potential role of L1 in mediating or increasing genome instability during tumorigenic development. Different host factors have been shown to influence L1 mobility through several mechanisms. However, systematic analyses of host factors affecting L1 retrotransposition are limited. Here, we developed a high-throughput microscopy-based retrotransposition assay and coupled it to a genome-wide siRNA knockdown screen to study the cellular regulators of L1 retrotransposition in human cells. We showed that L1 insertion frequency was stimulated by knockdown of Double-Stranded Break (DSB) repair factors that are active in the S/G2 phase of the cell cycle including Homologous Recombination (HR), Fanconi Anemia (FA) and, to a less extent, microhomology-mediated end-joining (MMEJ) factors. In particular, we show that BRCA1, an E3 ubiquitin ligase with a key role in several DNA repair pathways, plays multiple roles in regulating L1; BRCA1 knockdown directly affects L1 retrotransposition frequency and structure and also plays a role in controlling L1 ORF2 protein translation through L1 mRNA binding. These results suggest the existence of a “battle” between HR factors and L1 retrotransposons, revealing a potential role for L1 in development of tumors characterized by BRCA1 and HR repair deficiencies.

## Introduction

Retrotransposons replicate through RNA intermediates that are reverse transcribed and inserted at new genomic loci. LINE-1 (Long Interspersed element 1 or L1), the only active autonomous non-LTR transposable element in human cells, constitutes ~17% of the human genome and has had a substantial impact in the evolution of mammals^1–3^. Despite the fact that numerous copies of L1 are present in the human genome, most of them are 5’ truncated, rearranged or mutated and are therefore incapable of moving to new genomic locations through retrotransposition. Only about 100 L1 copies are potentially active in humans^3,4^ and only an handful of copies (dubbed “hot L1s”) have been shown to produce the majority of L1 activity^4^. The canonical, fulllength L1 element is ~6 kb long and consists of a 5’ UTR containing an internal promoter, two open reading frames (ORF1 and ORF2) and a 3’ UTR. ORF1p encodes a RNA-binding protein and exhibits chaperone activities^5^; ORF2p encodes a protein with endonuclease and reverse transcriptase activities^6,7^ The mobility of the L1 retrotransposon is fully dependent on its transcription and the subsequent translation of its encoded proteins, therefore both cytoplasmic and nuclear events are critical for L1 retrotransposition. In the cytoplasm, ORF1p and ORF2p bind to the mRNA that encoded them through unknown mechanisms that mediate cis-preference^8,9^ Interestingly, the translation of the second open reading frame of L1, ORF2p, is highly regulated in the cytoplasm of human cells. A still understudied and largely unknown unconventional termination/re-initiation mechanism has been proposed to explain the regulation of ORF2p translation^10^. Many trimers of ORF1p, as few as one ORF2p and the L1 mRNA form a LINE-1 ribonucleoprotein (RNP) complex which is thought to be an essential biochemical intermediate in mouse and human L1 retrotransposition^8,11^.

L1 RNP complexes are able to enter the nucleus mainly during mitosis^12^, and based on timing experiments, the subsequent steps in retrotransposition are thought to occur mainly during the S phase. The nuclear L1 mRNA is reverse transcribed and inserted in a new genomic locus through a partially understood molecular mechanism known as target-primed reverse transcription (TPRT)^13^. According to the largely accepted TPRT model the ORF2p encoded endonuclease domain makes a DNA nick on one strand of the host genome at AT rich consensus sequence (5’-TTTTAA-3’ consensus)^7,14–17^. The resulting 3’-OH end is extended by the reverse transcriptase domain of ORF2p that makes a DNA copy from the L1 RNA used as a template^6^. Cleavage of the second target genomic DNA strand usually occurs 7-20nt downstream of the first cut through an incompletely elucidated mechanism that results in the duplication of the DNA sequences between the two nicks (target site duplication or TSD)^18^. The entire TPRT process is completed by formation of a new double-stranded DNA copy of L1, flanked by target site duplications (TSDs)^19^. In addition to this canonical pathway, a second, less frequent L1 integration mode, independent of L1-endonuclease activity, was described both in a cell free system and in cells^6,20–22^. According to the endonuclease-independent insertion model, reverse transcription can be initiated from pre-existing chromosomal DNA breaks, for example at telomeres^21^. The majority of this type of L1 insertions have been measured at detectable frequencies in cell lines lacking non-homologous end joining (NHEJ) factors, facilitating a DSB repair mechanism which ligates together broken DNA ends, can function throughout all phases of the cell cycle, and is the predominant repair pathway in many mammalian cells^23,24^.

Due to its ability to move within the human genome, L1 retrotransposon has the potential to “jump” into coding regions and affect gene expression. Moreover, the repetitive nature of L1 in the genome was shown to increase chromosomal rearrangements, translocations and imprecise pairing during mitosis and meiosis^25,26^. About one hundred specific retrotransposition events have been documented to cause gene mutations leading to diseases in individual humans^27^. L1 retrotransposition has been also shown to induce genomic instability by generating DNA breaks^28–31^, a common feature in tumor initiating cells. More recent studies also demonstrated a wider prevalence of L1 expression in somatic cells, namely in neuronal stem cells, aging cells and different types of cancers^32^. Despite the many reports of L1 expression in somatic cells, very little is known about L1s’ physiological and pathological role during development or in the initiation and development of cancer.

DNA repair factors play essential roles in maintaining genome integrity; conversely, cancer risk and progression is closely linked to deficiencies in DNA repair pathways^33,34^. Several DNA repair factors have been shown to affect L1 retrotransposition but these observations are often inconsistent between studies, and sometimes the observed effects are quite modest. For instance, two groups have reported contradicting results on the role of ATM in regulating L1 retrotransposition, reporting either an activating^31^ or inhibitory^35^ role of ATM in L1 retrotransposition. In another case, NHEJ factors (Ku70, DNA-PKcs, Artemis, and Ligase IV) were also implicated as host factors (albeit in chicken cells, not the native host)^36^ while increased endonuclease-independent retrotransposition of LINE-1 was reported in XRCC4 and DNA-PKcs mutant cells^20^. Most recently, HR, Fanconi Anemia (FA) and Nucleotide Excision Repair (NER) proteins have been reported to play a role in suppressing L1 retrotransposition^24,37,38^. Despite the many studies, the relationship between DNA repair and L1 retrotransposition is still under continuous debate. It remains unclear, for instance, whether retrotransposition events facilitate the cellular DNA repair process or if the LINE-1 machinery simply takes advantage of potential priming sites offered by such breaks. Moreover, a molecular model that mechanistically explains the role of specific DNA damage repair factors on L1 retrotransposition is still lacking.

In order to gain a more systematic understanding of the cellular regulators of L1 retrotransposition, we undertook an arrayed, genome-wide siRNA knockdown screen utilizing a high-throughput microscopy-based approach. This screen uncovered hundreds of putative L1 regulators confirming and expanding a recently published CRISPR-based screen also set to identify regulators of L1 activity^38^. A clear result from our screen was that HR factors, and in particular the BRCA1 protein, had strong inhibitory effects on L1 retrotransposition. We provide evidence that BRCA1-mediated suppression occurs at multiple steps of L1 lifecycle: 1) directly during TPRT in the nucleus and 2) indirectly during L1 protein translation in the cytoplasm. Our results suggest that BRCA1-mediated HR (Homologous Recombination) repair competes with the L1 insertional process at pre-existing or at L1-created DNA breaks, and that resected DNA ends may represent a potential structure that is incompatible with completion of the TPRT reaction. In the cytoplasm, BRCA1 suppresses ORF2 translation through direct association with its mRNA. This leaves only the G1 phase, during which BRCA1 transcripts are less abundant, as a brief time window for ORF2p translation. These findings pave the way to the better understanding of L1 regulation in normal human cells as well as in cancers often characterized by the deficiency of DNA repair factors such as BRCA1 and Fanconi anemia proteins.

## Results

### Whole-genome siRNA screen using image-based retrotransposition assays

In order to study cellular factors that regulate L1 retrotransposition, we developed and optimized a high-throughput microscopy-based and relatively rapid retrotransposition assay in HeLa M2 cells^39^ (which constitutively express the rtTA or “Tet-ON” system) that allows measurement of L1 retrotransposition in an array-based format. This approach provides a more robust readout within a few days, and a less labor-intensive experimental procedure compared to traditional flow cytometry or selection-based retrotransposition assays. A synthetic L1 (ORFeus) retrotransposition reporter was expressed from a Doxycycline-inducible (Tet) promoter with its 3’UTR bearing an antisense GFP disrupted by an inverted intron (GFP-AI). This reporter design, extensively evaluated and used in previous studies^40,41^, allows quantification of retrotransposition by measuring the percentage of GFP positive cells (**Figure 1A**). We cloned the L1 reporter in the pCEP-puro vector for episomal expression, thus allowing the maintenance of a “quasi-stable” cell line under puromycin selection^40^. Cells carrying the L1 reporter were plated on 384 well plates with arrayed siRNA pools to knock down each individual human gene. 3 days after knock-down, the cells were stained with DAPI and the total number of cells and the number of green cells quantified by high content microscopy (**Figure 1A**). Measurement of retrotransposition was done after 3 d of L1 expression, because, in our setting, retrotransposition rate reaches saturation at this time point (**Figure S1A**). To ensure that L1 retrotransposition did not have a toxic effect on cells and to ensure that fluorescence measurements were unaffected by increased auto-fluorescence of dead cells, we also labeled dead cells after inducing expression of the L1 reporter for 3 d. Cells that underwent retrotransposition (GFP+ cells) were not stained in red (dead cells) and no bias towards dead cells was observed during automated tracking of green cells (**Figure S1B**) (See Methods). DAPI viability measurements were included in the whole genome screen and siRNAs that displayed excessive cell toxicity (less than 1500 cells left in the well before GFP and DAPI quantification) were discarded from the analysis. We also compared the distribution of z-scores from siRNAs resulting in low viability with the distribution of z-scores from the whole siRNA dataset. We noticed that siRNAs that induce cell death are significantly enriched in the class of knock down treatments that decrease L1 retrotransposition (“supporters”) (P<2.2e-16, Fisher’s Exact test) (**Figure S1C**).

**Figure 1.**
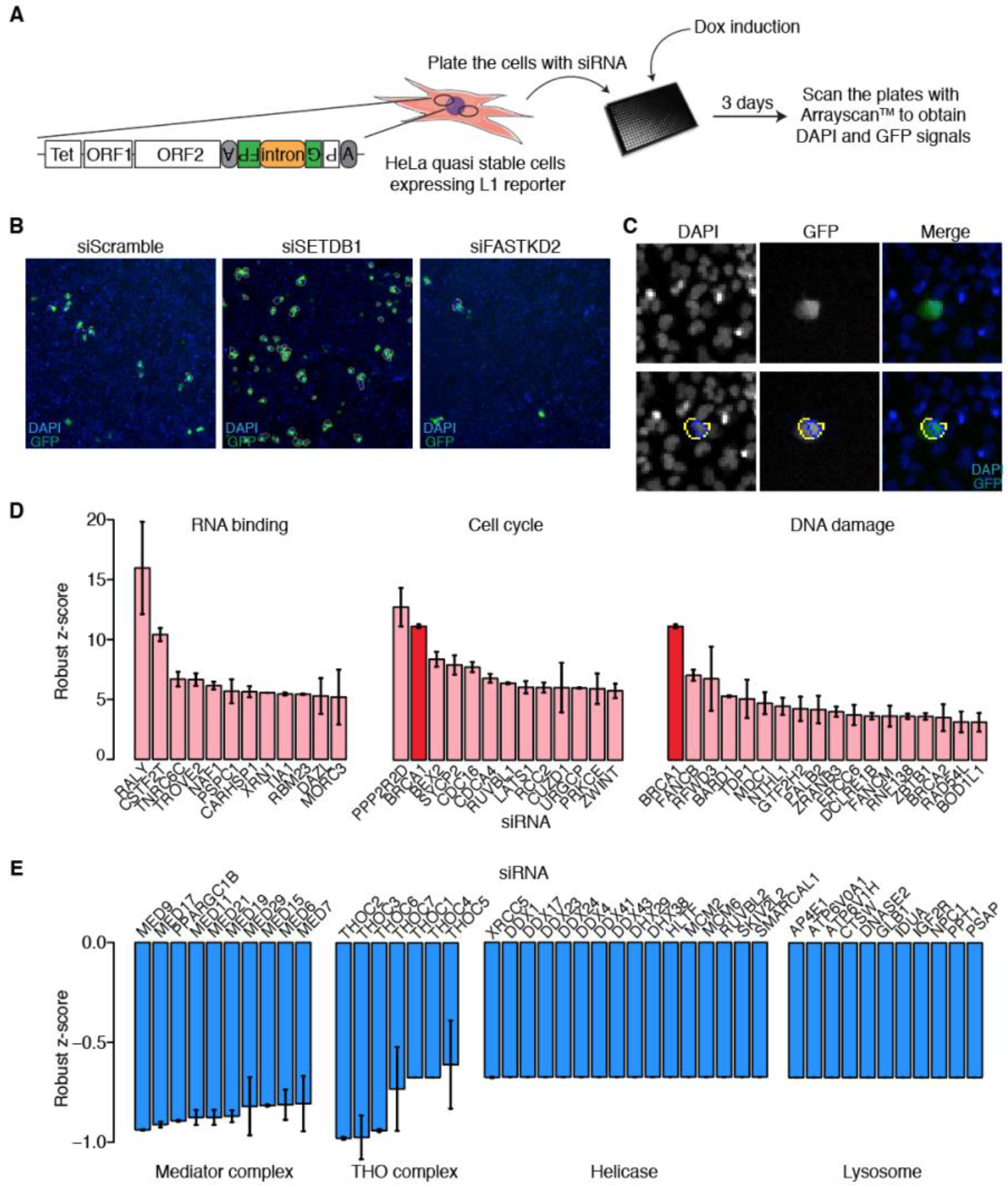
A whole-genome siRNA knockdown screen for L1 regulators. **A.** The workflow of the screen is presented. The L1 retrotransposition reporter includes ORFeus-L1 driven by a Tet-inducible promoter with an antisense intron-containing GFP engineered into its 3’UTR. This reporter is episomally maintained in HeLa-M2 cells cultured under puromycin selection. Cells expressing the L1 GFP-AI reporter were plated into 384-well plates containing arrayed pools of 3 siRNAs targeting the same human transcript. Doxycycline was added after 24 h to induce L1 expression. 3 days after induction, cells were fixed, permeabilized, stained with DAPI and scanned using the Arrayscan^TM^ high content microscope to quantify the GFP and DAPI signals. **B.** Examples of images obtained with Arrayscan™. 4 pictures are taken for each well at 5x magnification. siScramble knockdown represents the baseline L1 retrotransposition rate. Positive (e.g. siSETDB1) and negative (e.g. siFASTKD2) controls are shown as examples. **C.** Images were quantified using the Arrayscan™ built-in software. DAPI was used to detect and identify each nucleus (blue circle around cells); GFP signal (yellow circle around cell) was subsequently measured for each cell. **D.** Three top cluster of genes identified as L1 repressors from the primary screen are reported. *BRCA1* is identified in deep red bars. **E.** The four top clusters of genes identified as L1 activators are reported.

We performed a whole genome screen following the settings described above and using HeLa-M2 cells episomally expressing L1 (ORFeus) GFP-AI (**Figure 1A, Table S1**). Each plate included siRNA controls such as siRNA Scramble, siRNA against SETDB1 and siRNA against FASTKD2; SETDB1 was identified in a small-scale screen as a factor that, if knocked down, robustly and reproducibly enhanced L1 retrotransposition (referred to as a “inhibitor” of L1 retrotransposition) while FASTKD2 was identified as a factor that, if depleted, induced inhibition of L1 retrotransposition (referred to as a “supporter” of L1 retrotransposition) (**Figure 1B**). DAPI and GFP signals were quantified algorithmically using optimized parameters (**Figure 1C**). siRNAs displaying discordant results among the three replicates were discarded (red dots in **Figure S3B**). Upon screening and analysis, we identified 790 inhibitors (factors that if depleted increase L1 retrotransposition) and 1583 supporters (factors that if depleted decrease L1 retrotransposition) based on their robust Z-scores. GO term analysis of the inhibitors showed significant enrichment of genes involved in RNA binding, cell cycle and DNA repair (**Figure 1D**); whereas “supporters” of L1 retrotransposition significantly clustered in GO classes such as the mediator complex, THO complex, helicase and lysosome (**Figure 1E**). We noticed that several genes such as *BRCA1, BRCA2* and genes encoding components of the Fanconi Anemia pathway, involved in DNA damage repair and more specifically in double strand break (DSB) repair, strongly inhibited L1 retrotransposition. We therefore set up to validate and investigate the relevance of these group of genes on L1 activity.

### Secondary validations of DNA repair genes

Among the strongest hits established from our screen, we decided to focus on the DNA damage repair cluster (**Figure 1D**), a cluster also identified by^38^ using a CRISPR based screen, by Garcia Perez et al (cosubmitted) using a candidate gene approach and by Ardeljan et al (cosubmitted) who paradoxically identified this cluster of genes as synthetic lethal in p53^-^ L1-expressing cells. In order to understand which specific DSB repair pathways restrict L1 retrotransposition, we selected all 176 genes with the following GO term annotations: Fanconi Anemia (FA), Homologous recombination (HR), non-homologous end joining (NHEJ) and micro-homology mediated end joining (MMEJ). We then performed siRNA knockdown of these 176 selected genes in a 96 well format (**Figure 2A, Table S2**). This setting also allowed us to additionally perform an exon junction RT-qPCR assay to detect intronless GFP (from cells that had undergone retrotransposition) as an additional validation of retrotransposition at the genomic level; this assay should also, in theory, be independent of cell death (**Figure S2A**). We observed a good correlation between the results obtained in the 96 well setting versus the results in our whole genome screen (Pearson’s *r* = 0.57) with, as expected, a significant decrease in noise in the 96 well format (**Figure S3A, S3B**).

**Figure 2.**
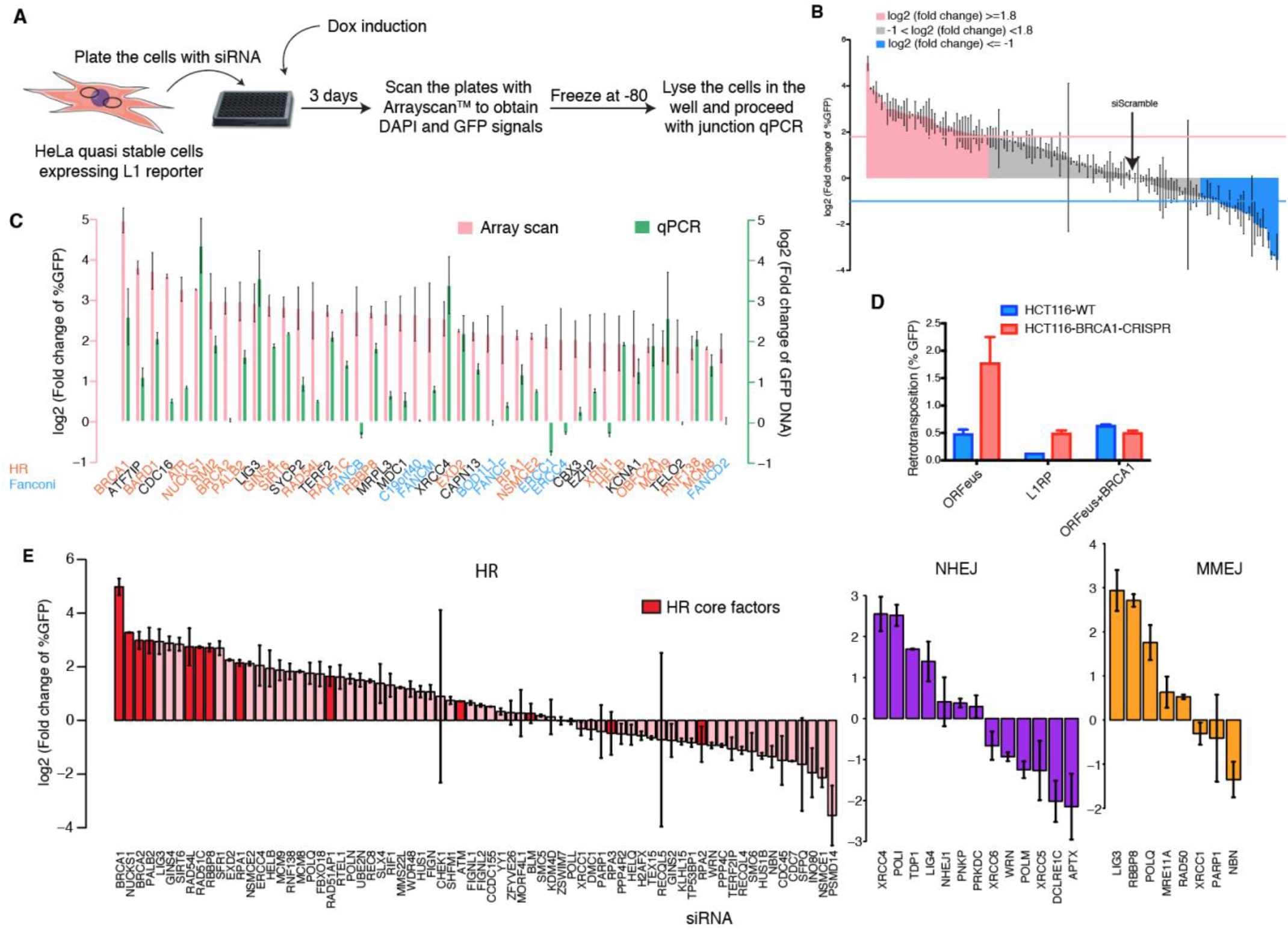
Secondary validations of the DNA repair factors. **A.** The workflow of the secondary validation is presented. siRNAs against DNA repair factors (whether or not they were revealed in the primary screen) were validated in 96-well format. The same cells were processed and analyzed for Arrayscan^TM^ analysis followed by GFP-AI and exon-junction qPCR (see text) using total DNA from each well. **B.** Fold changes (log2 scale) compared to siScramble control are plotted for each gene. Horizontal lines indicate chosen thresholds to define L1 retrotransposition repressors and activators (>=1.8 for repressors and <=-1 for activators). **C.** Exon-junction qPCR analysis is reported in green bars together with Arrayscan™ analysis in red bars for L1 repressors and blue bars for L1 activators. **D.** *BRCA1* was knockout in HCT116 cells infecting cells with lentivirus particles expressing CRISPR-Cas9 targeting BRCA1. These cells were used to preform rescue experiments by also transfecting a *BRCA1* expressing vector and quantifying retrotransposition (% GFP positive cells). GFP-AI reporters expressing recoded L1 (ORFeus) or non-recoded L1 (L1rp) were used. The expressed plasmids are indicated on the X axis, blue bars indicate wildtype HCT116 cells, red bars indicate *BRCA1* deficient HCT116 cells. **E.** Factors from the three major DSB repair pathways (HR, NHEJ, MMEJ) are grouped and their effect on L1 retrotransposition shown in a bar plot. The red bars indicate HR core factors.

Our GFP-AI exon-junction qPCR measurements, specific for spliced (retrotransposed) GFP DNA is in overall agreement with the results obtained in the screen and in our 96 well validation (**Figure S2B**). Indeed, most of the siRNAs with increased GFP signal also increased the spliced GFP RT-qPCR signal, and vice versa. Many siRNAs though, especially those that did not significantly change L1 retrotransposition (**Figure S2B**) showed discordant effects in these two measurements. Considering both measurements (96-well secondary screen and exon junction qPCR) for the siRNAs selected for secondary validations, we set a threshold of >1.8 (log2 fold change of the percentage of GFP+ cells) for inhibitors and −1 (log2 fold change of the percentage of GFP+ cells) for supporters (**Figure 2B**). 42 genes behaved as strong inhibitors of L1 retrotransposition and functionally clustered in the DNA repair GO class (**Figure 2C**).

### BRCA1-dependent HR pathway restricts L1 retrotransposition through resection

Among the inhibitors, BRCA1 showed the highest restriction of L1 retrotransposition. Corroborating the idea of an important role of BRCA1 and the HR repair pathway in L1 retrotransposition, we also identified members of the FA/BRCA1 pathway as strong inhibitors of L1 retrotransposition **(Figure 2C**, blue). Moreover, overexpression of BRCA1 was able to restore the L1 retrotransposition frequency to wild-type level in BRCA1 knockout cells (**Figure 2D**) in line with a direct role of BRCA1 in L1 retrotransposition repression. To understand how different DSB repair pathways respond to L1 activity, we analyzed genes in the three main DSB repair pathways defined by GO terms: HR, NHEJ and MMEJ. Among the genes characterized by HR GO term, we considered a subset as core DNA repair factors as defined by the KEGG pathway database and^42,43^ (Figure 2E, red bars). Both HR core proteins and MMEJ had a bias towards factors that inhibit L1 retrotransposition whereas NHEJ factors had a more ambiguous behavior (**Figure 2E**).

One of the common features of HR and MMEJ is that they both require resected DNA end structures. We therefore looked at RPA, Mre11 and Exo1 proteins, that are involved in the resection process, and noticed that, in our screen, all those proteins too functioned as L1 inhibitors (**Figure 2E**). This led us to hypothesize that DNA end resection (perhaps in the form of RPA bound ssDNA) may act as a physical barrier to L1 retrotransposition. To investigate whether BRCA1-dependent resection inhibits L1 retrotransposition, we performed epistasis analysis by knocking down resection factors in addition to BRCA1. We found that another key factor mediating resection, CtIP, also behaved as an L1 inhibitor, whereas 53BP1, shown to prevent end resection, acted as a L1 supporter. Lastly, we observed that the increased level of retrotransposition measured upon BRCA1 depletion, returned to basal level upon knocking down both 53BP1 and BRCA1 (**Figure 3A**). This result is in line with several studies showing that 53BP1 loss partially restores HR response in BRCA1-deficient cells by rescuing end-resection^44,45^. Similar effects on retrotransposition were observed using L1rp (the native L1 sequence) instead of recoded L1 (ORFeus) (**Figure S4**), indicating that the underlying BRCA1 mediated mechanism that restricts L1 activity is not RNA sequence dependent.

**Figure 3.**
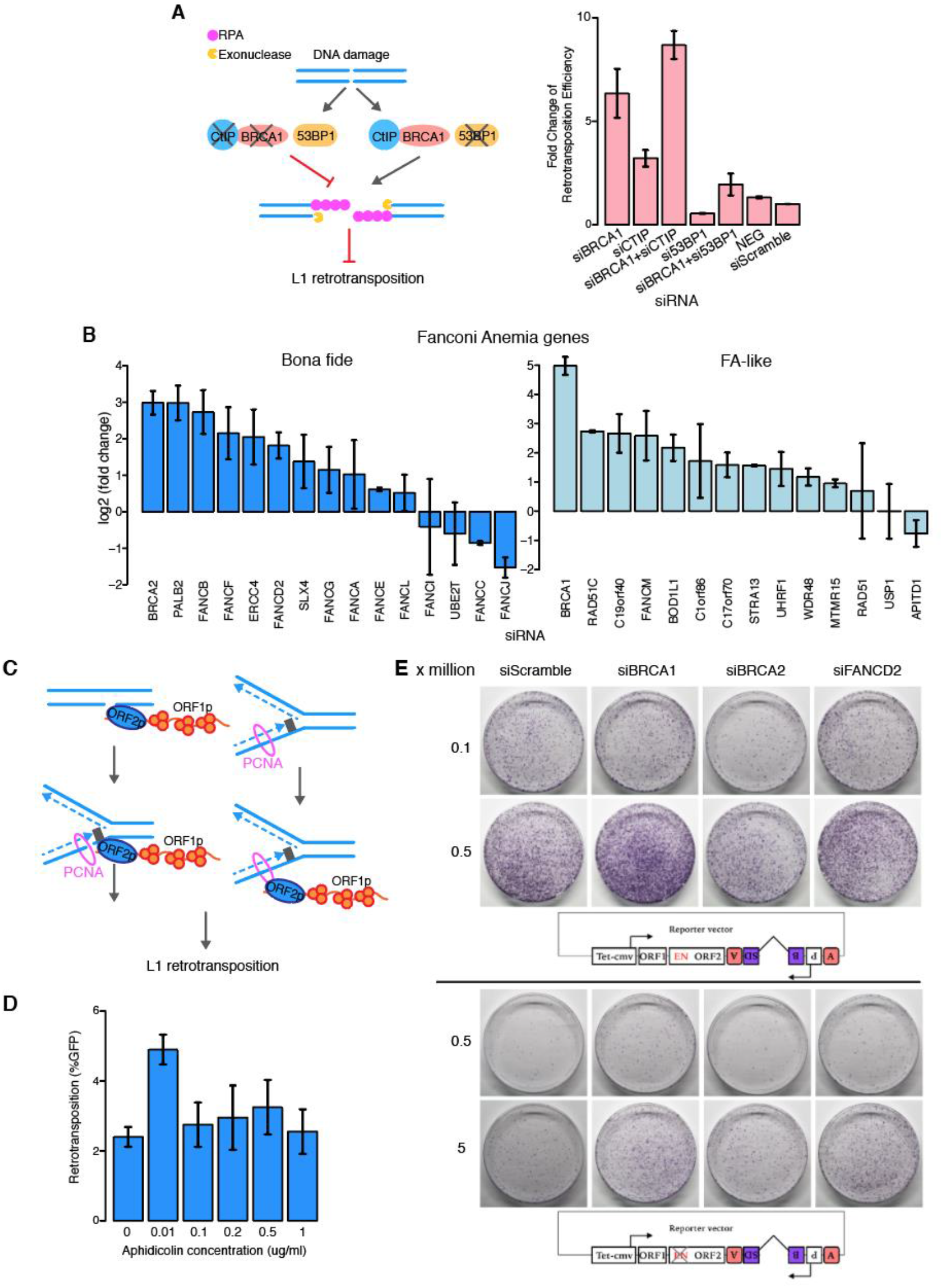
HR/FA pathways strongly repress L1 retrotransposition. **A.** Epistasis analysis was performed to assess the effect of DNA end resection on L1 retrotransposition. left panel: *BRCA1* and *CtIP* knockdowns promote, whereas *53BP1* knockdown inhibits DNA resection around a DNA break. Resection can still occur when *BRCA1* and its antagonist *53BP1* are both absent. Right panel: single and double knockdown of the indicated proteins was performed using HeLa cells plated in 6-well plates. Fold change of L1 retrotransposition was calculated by normalizing %GFP upon knockdown to %GFP of scrambled siRNA control treated cells. **B.** Canonical FA factors and FA-like factors are plotted for their effect on L1 retrotransposition. The factors within each group were ordered by their fold change compared to siScramble control. **C.** Upper panel: proposed model for L1 insertion occurring at stalled replication forks via ORF2-PCNA interaction. Bottom panel: L1 retrotransposition is measured in cells treated with increasing concentrations of Aphidicolin, to induce slowing or stalling of replication progression. **D**. Retrotransposition assays using a BSD-AI reporter. Clones of cells after Blasticidin selection are stained with crystal violet and presented. The number of cells plated per well and the transfected siRNA are indicated. Schematic representation of each of the L1-BSD-AI reporter plasmid used in the experiment are shown on the bottom.

### Fanconi Anemia factors limit L1 retrotransposition during replication

Our whole-genome screen also identified a cluster of L1 inhibitors within the FA pathway^46^, which acts to recognize and repair lesions at stalled replication forks. Interestingly, most FA factors behave as inhibitors of L1 activity except for FANCJ, which acts as a strong supporter (**Figure 3B**). Because of the central role of FA factors in protecting replication forks arrested by DNA damage sites^47^ we focused on better elucidating the role of L1 retrotransposition in potentially creating and/or exploiting stalled replication forks. To investigate whether L1 insertions occur at stalled replication forks or alternatively, whether L1 itself creates stalled forks e.g. upon endonuclease cutting (**Figure 3C**), we measured the retrotransposition efficiency in cells treated with different concentrations of Aphidicolin. We observed that L1 insertions increased more than two-fold under low concentrations of Aphidicolin treatment and remained unaltered at higher concentrations (**Figure 3D**). As a low concentration of Aphidicolin slows down and stalls replication fork progression but does not arrest the cell cycle, this result points toward a possible recruitment of L1 to stalled DNA replication forks, where retrotransposition may occur at high frequency (see Discussion).

### BRCA1 depletion increases endonuclease dependent and independent L1 retrotransposition

It has been previously shown that L1 is also able to retrotranspose at pre-existing DNA damage sites and that these events are independent of L1 endonuclease (EN) enzymatic activity^20,21^. Because BRCA1 depletion causes impaired DNA damage repair function and therefore an increase in the number of unresolved DNA damage sites^48,49^, we hypothesize that, at least in part, the increase of retrotransposition observed in the absence of BRCA1 may be due to EN-independent activity of L1. Due to the rarity of endonuclease-independent retrotransposition events, we utilized a retrotransposition assay based on drug selection (Blasticidin resistance selected over 10 days) instead of GFP fluorescence detection, to quantify L1 insertions. As expected from our previous results, BRCA1 depletion as well as depletion of a centrally important FANC protein (FANCD2) induced an increase of L1 EN-dependent retrotransposition (**Figure 3E**, upper panel). Utilizing an endonuclease dead L1 BSD-AI reporter (**Figure 3E**, bottom panel), we also found that knock down of BRCA1, BRCA2 and FANCD2 increased EN-independent retrotransposition, demonstrating a role of the HR/FANC pathway on both EN-dependent and – independent retrotransposition (**Figure 3E**).

Surprisingly, and in contrast with our measurements based on the GFP-AI reporter, we observed that knock down of BRCA2 induced a paradoxical decrease in colony formation upon L1 EN-dependent retrotransposition (**Figure 3E**, upper panel, compare siScramble vs siBRCA2). This effect of BRCA2 knock down was not observed using an EN mutant L1 reporter (**Figure 3E**, bottom panel, compare siScramble vs siBRCA2). This observation supports a possible longer-term synthetic fitness interaction between L1 overexpression and depletion of BRCA2, as also observed by Ardeljan et al. (cosubmitted) (see Discussion). Moreover, using the BSD-AI reporter we observed a smaller effect of BRCA1 knockdown on L1 retrotransposition in this longer-term assay, compared to the three-day readout employed with GFP-AI (**Figure 3E and 1D**). As for our observations on BRCA2 knock down, the effect of BRCA1 knockdown was possibly reduced in BSD-AI experiments due to cell death during a longer culture period. In the case of BRCA2 knockdown, the number of retrotransposed clones even decreased compared to control, however, the opposite was observed in the absence of L1 endonuclease activity (**Figure 3E**). These observations suggest that elevated L1 retrotransposition is detrimental to the long-term growth of cells deficient in HR/FA pathways.

### BRCA1 depletion induces formation of target site deletions upon L1 retrotransposition

In order to understand the molecular mechanism of BRCA1-mediated suppression on L1 TPRT (target primed reverse transcription) specifically, we sequenced the *de novo* L1 insertions and their flanking regions in the cells depleted of BRCA1, using a modified recovery assay^20^ (**Figure 4A**). Several features of the L1 retrotransposition sites were evaluated, including the sequence at which the first nick occurred (target site), the length of the inserted polyA, the length of the L1 insertions and the type of L1 insertions (inversion at 5’ end versus canonical) (**Table S3**). No significant differences were observed in the length of inserted L1 or polyA comparing control cells to cells depleted of BRCA1 (**Figure S5**). Moreover, we observed that the majority of L1 insertions are characterized by a recognizable L1 A/T rich target site (TTTT AA) demonstrating that most L1 insertions were mediated by ORF2 endonuclease activity. To support this result, we also quantified the distances of L1 insertions to the common fragile sites (CFSs). During DNA replication, CFSs constitute “difficult sites” at which increased DNA breaks have been measured^50,51^. As expected, L1 insertions were not enriched at these CFSs, suggesting that most L1 insertions that happened in BRCA1/FA deficient cells were endonuclease-dependent insertions (**Figure 4B**).

**Figure 4.**
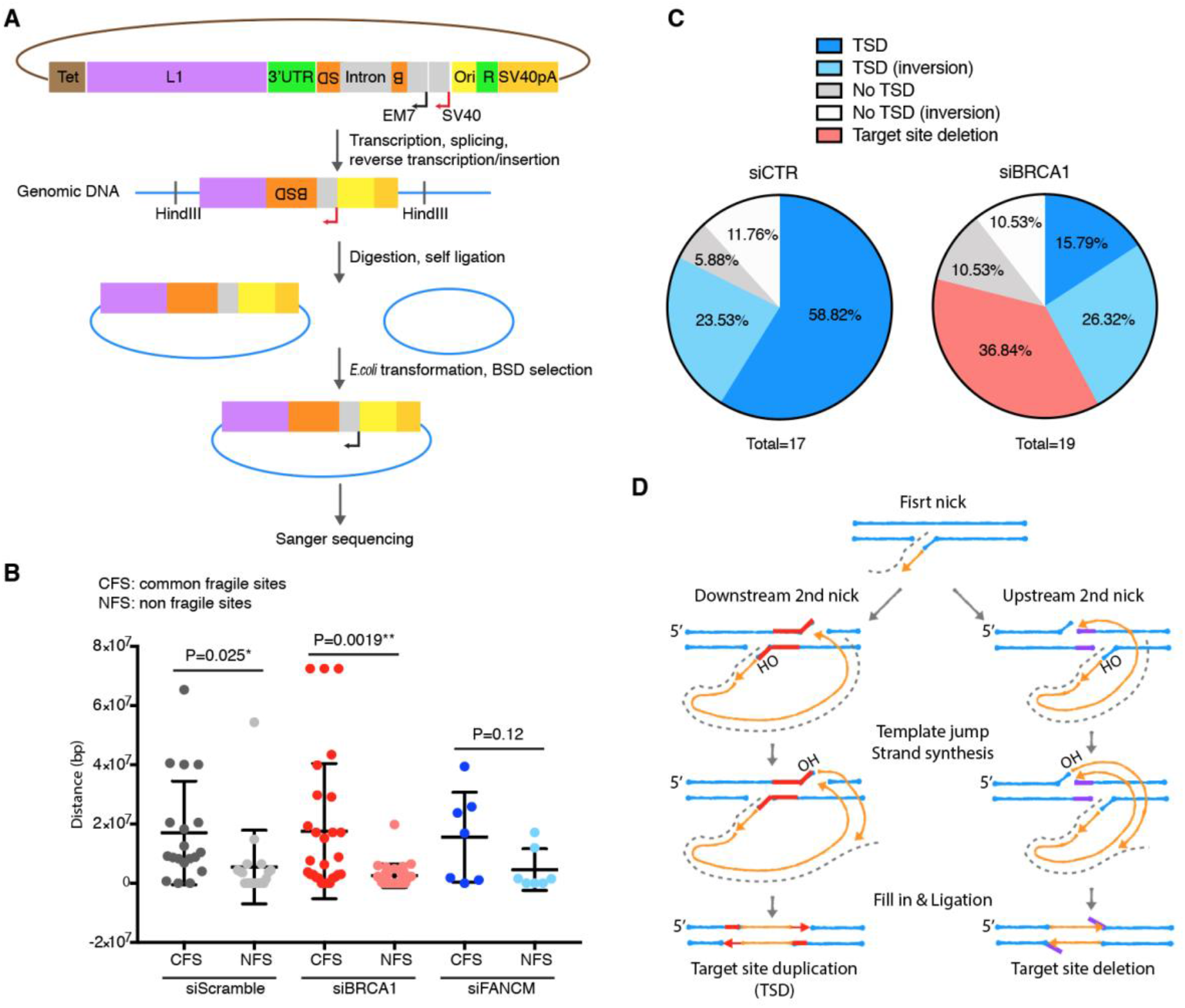
BRCA1 deficiency induces increased formation of target site deletions upon L1 retrotransposition. **A.** Diagram depicting the procedure used to recover and analyze *de novo* L1 insertions. A BSD-AI cassette was introduced into the 3’UTR of L1 together with the EM7 bacterial promoter (gray box also including the SV40 promoter) and a bacterial DNA origin of replication (yellow box). The Blasticidin is expressed from a mammalian promoter (SV40) after a retrotransposition event occurs. Genomic DNA of Blasticidin resistant clones were then extracted, subjected to HindIII restriction digestion, self-ligated and transformed into bacteria. Blasticidin resistant bacteria colonies were recovered and sequenced. **B.** Distance of the *de novo* L1 insertions to common fragile sites (CFS) or non-fragile sites (NFS). **C.** Pie charts showing the percentages of the different L1 insertion types detected in the control (scrambled siRNA) and BRCA1 deficient cells (siBRCA1). **D.** Proposed model for target site deletions in BRCA1 deficient cells. Genomic DNA is shown in blue; L1 mRNA is shown as dashed lines and L1 cDNA is shown in orange. Sequences between the two nicks induced by L1 are shown in red for target site duplications (TSD) and purple for target site deletions.

We observed a striking increase in the formation of target site deletions relative to target site duplications upon BRCA1 knockdown. (**Figure 4C**). L1 insertions causing target site deletions have been previously observed in the human and Drosophila genome^17,52,53^. The major determining factor between target site deletions and the more common and mostly observed target site duplications is likely to be the position of the second DNA nick relative to the first bottom strand nick produced during TPRT. During canonical L1 TPRT, the second nick is usually “downstream” of the first nick, whereas for target site deletion, the second DNA nick is inferred to occur upstream of the first nick (**Figure 4D**). The reason for the biased preference for “downstream” second nicking during L1 TPRT is not known, but BRCA1 seems to be a strong determinant of this bias.

### BRCA1 inhibits ORF2 translation in G1 phase of the cell cycle

During our study of L1 in BRCA1-depleted cells, we were surprised to see evidence for an additional distinct layer of regulation of L1 induced by BRCA1. We observed that BRCA1 depletion induced a specific increase in L1 ORF2p for both ORFeus and L1rp (non-recoded L1) while ORF1p level was not affected (**Figure 5A, S6**). Because ORF1p and ORF2p are translated from a bicistronic mRNA, we hypothesized an effect of BRCA1 depletion on ORF2p translation. Immunoblot analyses of cells depleted of BRCA1, 53BP1 and FANCM proteins, revealed that ORF2p levels increased in BRCA1-depleted cells while ORF1p remained unaltered. Interestingly, this translational effect was independent of the HR repair pathway as knocking down BRCA1 together with 53BP1 did not rescue this increase of ORF2p (**Figure 5A**). This effect was also validated using immunofluorescence staining which showed a clear increase in ORF2p signal upon knocking down BRCA1 compared to control siRNA treated cells (**Figure 5B**). BRCA1 expression changes during the cell cycle decreasing in G1 and increasing in S, G2 and M phase (cyclebase 3.0). To verify the cell-cycle dependent expression of BRCA1 we sorted HeLa S.FUCCI cells in G1 phase (cells with red nuclei) from cells in the rest of the cell cycle (cells with green nuclei). As expected, cells in G1 phase express less BRCA1 protein while cells in S/G2/M express more BRCA1 compared to an unsorted population (**Figure 5D**, left panel). In line with the inhibitory effect of BRCA1 on ORF2p translation (**Figure 5A**), we also observed that cells expressing low levels of BRCA1 protein (most likely in G1 phase) were more likely to express ORF2p, whereas cells displaying high levels of BRCA1p did not usually express ORF2p (**Figure 5C, 5D** right panel; p-value: 1.7 x 10^-9^).

**Figure 5.**
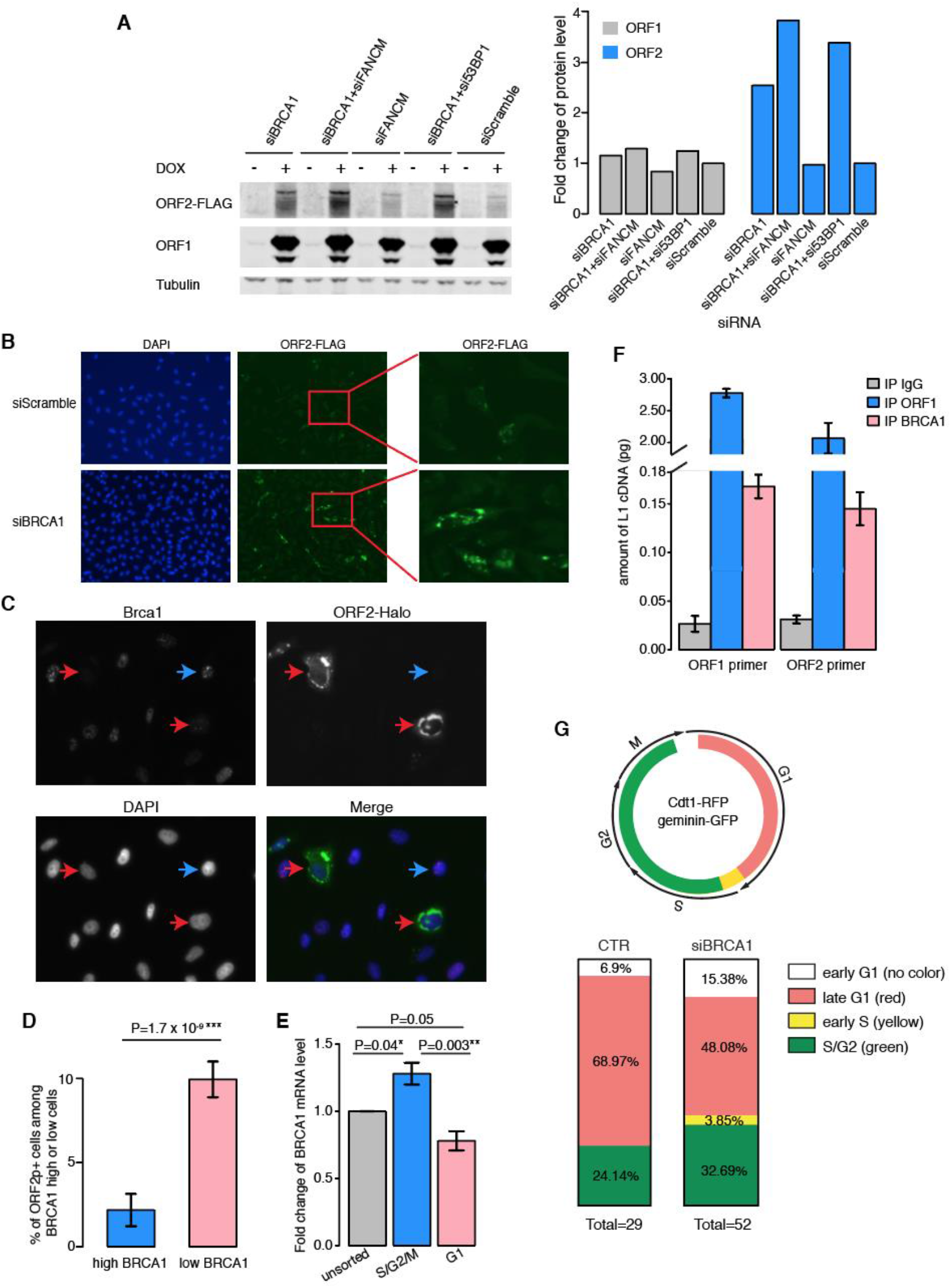
BRCA1 inhibits ORF2 translation in the cytoplasm. **A.** Left panel: Western blot of ORF1 and ORF2-FLAG proteins from control HeLa cells (siScramble) or cells treated with the indicated siRNAs and expressing L1 ORFeus (untagged ORF1p, Flag-tagged ORF2p). Doxycycline (DOX) is used to induce the expression of L1 under a Tet-regulated promoter and tubulin is used as loading control. Right panel: quantifications of ORF1p and ORF2p bands normalized by the corresponding signal in the siScramble control treatment set as 1. **B.** Immunofluorescence of HeLa cells expressing a recoded L1 with FLAG-tagged ORF2 and treated for 24 hrs with control siRNA (siScramble) or siRNA against BRCA1 (siBRCA1). **C.** HeLa cells expressing a recoded L1 with halo-tagged ORF2p detected with JF657 Halo-tag (green pseudocolor). BRCA1 protein was also detected (red pseudocolor) using a specific antibody (D-9). **D.** quantification of Figure 4C showing mutually exclusive patterns of BRCA1 and ORF2p. **E.** BRCA1 mRNA quantification in different phases of the cell cycle using sorted FUCCI cells. **F.** Amount of L1 mRNA bound to BRCA1. BRCA1 was immunoprecipitated from whole cell lysate from HeLa cells expressing recoded L1. L1 mRNA pulled down with BRCA1 was measured by qPCR and quantified compared to a standard curve of known concentrations of L1 DNA. G. Upper panel shows cell cycle stages indicated by colors in HeLa S.FUCCI cells. Bottom panel: cell cycle stages in which ORF2 protein started to be detected were quantified in control and *BRCA1* knockdown cells. Stacked bar plots showing the percentages of cells starting to express ORF2p in the different cell cycle stages.

We therefore hypothesized that BRCA1 may inhibit ORF2 translation through direct binding to L1 mRNA in the cytoplasm as such a function has been previously attributed to BRCA1 for several mRNAs and proteins^54^. To evaluate a possible physical interaction between BRCA1 and L1 mRNA, we performed immunoprecipitation of BRCA1 protein followed by RT-qPCR of L1 mRNA. We observed a >5 fold enrichment of L1 mRNA binding to BRCA1 compared to an IgG control (**Figure 5F**). Because BRCA1 expression is regulated during the cell cycle (**Figure 5E**), we measured the timing of ORF2 protein translation and expression during the cell cycle and interrogated the possible role of BRCA1 in this regulation. Using live cell imaging of HeLa S.FUCCI cells expressing an inducible ORF2-Halo tagged LINE-1^12^ we observed that ORF2 protein usually starts to be detectable during G1 phase, when BRCA1 transcripts are low (**Figure 5G, Movie 1**). This result clearly shows a cell-cycle dependent regulation of ORF2p translation. To determine the role of BRCA1 in this regulation we performed a similar analysis in cells depleted of BRCA1 by siRNA treatments. HeLa S.FUCCI expressing ORF2p-tagged L1 and depleted of BRCA1 showed an increased fraction of cells that express ORF2p in cell cycle stages other than G1 (**Figure 5G, Movie 2**) suggesting that BRCA1 inhibits ORF2p translation mainly in S/G2/M phases when BRCA1 is highly expressed. These data suggest that BRCA1 regulates L1 ORF2p translation through binding to its mRNA, possibly in a cell cycle regulated fashion.

Overall, our results demonstrate several lines of defense deployed by BRCA1 against L1 retrotransposon activity. In the nucleus BRCA1 surveils DNA breaks, promptly repairing them and rendering them inaccessible to potential L1 insertion. Moreover, BRCA1 directly affects L1 target primed reverse transcription (TPRT) inhibiting it and also preventing the formation of target site deletions most likely triggering DNA resection events able to inhibit L1 retrotransposition. Finally, BRCA1 is also able to repress L1 activity in the cytoplasm, where it inhibits L1 ORF2p translation through binding to L1 mRNA. These lines of regulation of L1 activity may be misregulated in certain BRCA1 deficient cancers highlighting a possible role of LINE-1 in the progression of tumors such as breast and ovarian tumors (see Discussion).

## Discussion

### Whole genome knockdown screen for regulators of L1 retrotransposition

In this study we conducted a whole-genome knockdown siRNA screen for the identification of factors affecting L1 retrotransposition. The setup of the screen was based on adapting to the higher throughput fluorescent-AI retrotransposition assay traditionally based on low throughput measurements to quantify retrotransposition events such as flow cytometry or colony formation assays. We utilized a microscopy-based method for the quantification of GFP fluorescence/retrotransposition, which extensively reduced experimental burden and increased the sensitivity of the assay compared to flow cytometric measurements. The entire experimental procedure takes less than one week without any need to replate or transfer cells. The increased performance of our microscopy-based assay compared to traditional assays is mainly due to:

1. High throughput output of the microscopy-based assay; the use of a high content microscope such as the Arrayscan VTI system simplifies the whole procedure to 3 steps: plating, induction and measurement.
2. The imaging approach is a bona-fide single cell analysis that is less affected by background auto-fluorescence and background signal compared to flow cytometry analysis.
3. Different from selection-based assays which require a long selection (10 days of culturing cells), the microscopy-based assay takes only 3 days from plating to measurement, minimizing technical variables introduced by long term siRNA treatments. An example of this phenomenon is the knock down of the *BRCA2* gene that performed as a clear L1 inhibitor in GFP-based experiments while it acted as a weak supporter using BSD-AI reporters (**Figure 3E**). We observed that *BRCA2* knockdown exerts a toxic effect on cell growth and longer cell culture led to the death of retrotransposition positive cells. This behavior was also observed using *BRCA1* siRNA in the presence of L1 activity although the toxic effect of *BRCA1* siRNAs had a much lower effect on cell growth compared to *BRCA2* siRNA. This observation may, to some extent, explain the low degree of overlap between hits from our screen and hits from the previously published CRISPR-based screen^38^.

Despite the advantages mentioned above, few limitations characterized our siRNA screen. First, the use of a Doxycycline-inducible promoter to drive the expression of L1 instead of the endogenous 5’UTR/promoter, restricts our analysis to factors affecting L1 activity post-transcriptionally. Any transcription factor that regulates L1 transcription or proteins that bind L1 5’UTR cannot be identified by our screen as L1 regulators. Also, the use of recoded ORFeus, which is characterized by higher L1 mRNA, protein and retrotransposition frequency, may have excluded the identification of factors that rely on native L1 sequence such as the HUSH complex^55^. On the other hand, the use of an inducible promoter gives the advantage of precise control of timing of L1 activation, avoiding possible long-term toxicity effects of L1 expression. Secondly, we observed that the hits in our screen are strongly biased towards L1 inhibitors versus supporters. This bias is probably explained by the fact that our baseline L1 retrotransposition frequency was around ~1%, and thus a further reduction of the percentage of retrotransposition events is at the boundary of statistically robust detection. To increase the assay sensitivity and confidence, we validated DNA repair genes based on GO term annotation with more cells (96-well plates) and by genomic exon-junction qPCR analysis of spliced GFP (**Figure S3**). Because of the discrepancies observed between the two methodologies (fluorescence measurement versus exon-junction qPCR analysis) we decided to use both methods to assess the effect of a particular factor on L1 retrotransposition. Our method is amenable to the high-throughput measurement of both GFP+ cell and spliced GFP signal from the same wells enabling faster acquisition of more reliable data.

### HR factors restrict L1 retrotransposition

Our screen clearly identified BRCA1 as a major inhibitor of L1 retrotransposition. Validation and characterization of this considerable effect of BRCA1 on L1 life-cycle revealed that BRCA1 plays several roles in restricting L1 function. Our data show that the increased L1 retrotransposition induced by BRCA1 depletion is mostly due to a direct effect on endonuclease-dependent retrotransposition and L1 TPRT. Indeed, sequencing of L1 insertions of cell depleted of BRCA1 showed that the majority of L1 insertions were characterized by a recognizable L1 endonuclease target site (TTTTAA). Furthermore, increased level of target site deletions in BRCA1 knockdown cells suggests that BRCA1 plays a role in determining the position of the second DNA nicking during TPRT, potentially directly blocking second nicking upstream of the fist nick. The bias toward “upstream” nicking observed in our experiments suggests a distinct physical DNA end structure in cells in which BRCA1 has been depleted. We nevertheless also observed that endonuclease-independent retrotransposition events were increased after knocking down *BRCA1* (but were not the major events inhibited by BRCA1), which indicates that, as previously shown^6,20,22^, unrepaired DNA breaks, increased by the lack of BRCA1 and other components of the HR DNA damage response pathway, can serve as potential targets of L1.

Because of our previous observations demonstrating that L1 retrotransposition is biased towards the S phase and that ORF2p binds components of the DNA replication fork (namely PCNA and MCM proteins)^12^, we hypothesize that BRCA1 and the homologous recombination (HR) pathway, the dominant pathway active during S/G2 phase of the cell cycle, competes with L1 retrotransposition. Consistent with this hypothesis, we observed that knockdown of components of HR and, to a lesser extent MMEJ, enhance L1 retrotransposition frequency, implying that these pathways interfere with retrotransposition. The mechanistic similarities between HR and MMEJ include the following features: 1) both pathways dominate during the S/G2 phase of the cell cycle^56^, the time at which L1 activity peaks in these cells^12^ 2) both pathways require resected DNA end structures to perform strand invasion, homology search and subsequent recombination. The above observations led us to hypothesize that DNA end resection complexes may act as a physical barrier to L1 retrotransposition, more specifically inhibiting the process of TPRT that requires a 3’OH as a priming template for reverse transcription. Moreover, the NHEJ DNA repair pathway acts in competition with the HR pathway; the choice between HR and NHEJ largely depends on the DNA end structures created by resection around the DNA damage site and is largely segregated within the cell cycle, with NHEJ highly active in G1 and HR much higher in S/G2^57,58^. Secondly, we observed that, during TPRT, in BRAC1 knockdown cells, L1 is seemingly able to choose between a downstream or upstream DNA second nick without bias, suggesting that in normal cells, BRCA1 and associated proteins may sterically block the formation of the second DNA nick upstream of the first nick during L1 retrotransposition (perhaps through the creation of resected end structures). It is possible that DNA repair factors normally accumulate upstream of the first nick, make it inaccessible to the EN domain of ORF2p.

These observation links up-regulation of L1 to cancers characterized by misregulation of the HR pathway such as mutations of BRCA1 and/or BRCA2 proteins, known to drive several cancers including breast and ovarian cancer. It is therefore possible that, in the context of misregulated HR, the expression of L1 and its activity may increase genome instability, accelerating tumor progression or the development of resistant clones upon treatment. Further studies are necessary to evaluate the role of L1 expression on tumor progression in tumors with a compromised HR background and to evaluate the possible use of L1 expression and activity as a biomarker for specific stages of tumor progression.

### Synthetic lethality and retrotransposition permissivity

Our results and those of^38^ are very consistent in that they indicate that retrotransposition activity is enhanced by knockdown or knockout of BRCA1 and FA factors. However, Ardeljan et al. (cosubmitted). paradoxically observe exactly the same sets of genes as synthetic lethal with LINE-1 overexpression. How can we resolve this apparent paradox? We note that our group and Liu et al. both used a relatively short-term readout of retrotransposition efficiency based on GFP (3 days in our case and 10 days for^38^) whereas Ardeljan et al. (cosubmitted). examined cultures after 27 days of selection for expression of a drug resistance marker. In all three cases, the experiments were done in p53 negative cells with retrotransposon overexpression. Thus, we rationalize this apparent contradiction as follows: knockdown/knockout of these genes has two effects: substantial increase in retrotransposition and long-term synthetic fitness effects. Consistent with this interpretation, when we examined the effect of *BRCA1* and *BRCA2* knockdown in a longer-term drug resistance-based selection, the enhancement of retrotransposition was far more modest and, in the case of *BRCA2* knock down, it even showed an opposite effect on retrotransposition compared to what was seen in the GFP readout (**Figure 3E**).

### L1 and the DNA replication fork

We imagine two possible interpretations of the connection between retrotransposon TPRT complexes and stalled replication forks (**Figure 3C**)^59^. One intriguing possibility is that L1 TPRT complexes create stalled forks, and that consequent recruitment of the FA complex leads to aggressive disassembly/repair of TPRT intermediates in addition to replication fork restart. In contrast, it may be that the retrotransposition complex actively targets stalled replication forks. Our recently published data^12^, supported by the results presented here, suggest that L1 may preferentially insert into the genome at stalled replication forks, as this genomic structure may provide an “open” environment most accessible to a large bulky retrotransposition complex. We are therefore more incline to the hypothesis that L1 targets stalled replication forks for TPRT rather than creating them. Previous study has shown that Tf1, a Long Terminal Repeat (LTR) retrotransposon, was guided to integrate at the stalled replication forks by the DNA binding protein Sap1^60^. In addition, recent work^61^ suggests that AT rich and T_n_|A_n_ motifs are determinants of replication origin usage in mammals and that they represent polar replication barriers where the replication fork tends to stall. Considering the genomic features of L1 insertions, these Tn|An tracts may 1) serve as canonical L1 retrotransposition target sites 2) be created by historical insertions of L1 itself, indicating a possible contribution of L1 in the evolution of replication origins. The data presented by^61^ seems to be in line with a potential recruitment and retrotransposition of L1 into stalled replication forks at T_n_|A_n_ sites where long unprotected single stranded stretches of As are created by the decoupling of the MCM helicases and DNA polymerase. We hypothesize that these regions are ideal sites for L1 retrotransposition.

FA proteins are known to be recruited at stalled replication fork^62^. Our results overall suggest that FA anemia proteins may inhibit retrotransposition of L1 during S phase, taking advantage of DNA replication fork stalling induced by damage. Treatment with low concentrations of aphidicolin and L1 retrotransposition measurements seem to support this hypothesis, however, the effect observed with aphidicolin treatment was only up to a 2-fold increase and thus seems insufficient to explain the significant increase observed when HR/FA factors pathways were impaired. We hypothesize that at these sites, L1 competes with the HR repair machinery for the use/repair of the nicked DNA. Once retrotransposition is initiated, L1 will be able to insert at least a partial L1 sequence in the new genomic locus; however, when BRCA1 and the HR machinery is recruited and is able to initiate DNA resection, L1 retrotransposition will be inhibited. Some questions including how L1 recognizes sites of stalled replication or whether L1 may be the cause of slowed replication forks remain unanswered. Moreover, we showed that L1 insertions were not enriched at CFS, indicating that the majority of the increased L1 insertions were not promoted by an elevated level of DNA breaks.

Future *in vivo* and *in vitro* studies of the molecular mechanisms of the role of BRCA1 on L1 retrotransposition may shed new light on the still largely uncharacterized process of L1 TPRT. It is still unclear whether the EN activity of ORF2p is responsible for both or just one of the two DNA nicks necessary for L1 retrotransposition. Also, a detailed model of L1 retrotransposition that incorporates the replication fork is still lacking. Our discoveries point towards a complex interaction between DNA replication fork, DNA repair machinery and L1 retrotransposon RNPs that leads to specific and well-regulated process culminating with what is considered a canonical L1 insertion normally observed all over the human genome. The better understanding of these pathways is a necessary step for the study of the role of L1 in cancer and the potential pharmacological manipulation of L1 retrotransposition in these tumorigenic settings characterized by misregulated DNA damage response. Interestingly, in tumors deficient of BRCA1/2, replication fork stability has been implicated in the development of cisplatin and PARP1 inhibitor resistance^63^ underlying the importance of better understanding the functional interactions between LINE-1, DNA replication and BRCA1.

### L1 ORF2p translation and BRCA1

Translational control of ORF2 remains a completely unknown process and here we observed an inhibitory effect on ORF2 protein level by BRCA1 in the cytoplasm. This effect was RNA sequence independent as the effect of BRCA1 depletion affect ORF2 level in both L1RP and ORFeus expressing cells. We therefore hypothesize that this inhibition may be dependent on L1 polyA tail. PABPC1, a L1 interactor, was previously shown to be essential for L1 RNP formation; it binds polyA tail and may potentially explain how BRCA1 can recognize and bind L1 mRNA. In addition, we also observed a cell cycle regulation of ORF2p translation that, in most cells, takes place during G1 phase. Because BRCA1 was previously shown to be regulated during the cells cycle we tested if BRCA1 repressed ORF2p translation in a cell cycle fashion. We observed a mild disruption on cell cycle control of ORF2 translation when BRCA1 is depleted, suggesting that BRCA1 repression of ORF2p translation may be linked to the cell cycle. The small amplitude of the effect induced by BRCA1 depletion may be due to incomplete depletion of BRCA1 in some of the observed cells or may be explained by the fact that BRCA1 is, most likely, not the only protein regulating ORF2p translation. Future experiments, possibly tracking both BRCA1 and ORF2p during various stages of the cell cycle will need to be designed to better elucidate the role of the BRCA1 pathway on ORF2p translation in cycling cells.

## Methods

### Cell Culture and Cell lines

HeLa M2 cells (a gift from Gerald Schumann, Paul-Ehrlich-Institute) were cultured in DMEM media supplemented with 10% FBS (Gemini, prod. number 100–106) and 1 mM L-glutamine (ThermoFisher/Life Technologies, prod. number 25030–081). HCT116+ch3 cells (an extra copy of chromosome 3 corrects a mismatch repair deficiency) were cultured in DMEM/F12, HEPES (ThermoFisher Scientific, catalog number 11330032) supplemented with 10% FBS and 400 μg/ml G418.

### Plasmids and DNA constructs

pLD401 (ORFeus with 3xFlag ORF2) and pMT302 (L1rp with 3xFlag ORF2 and L1 5’UTR) plasmids were previously described and characterized in^40^. pEA79 (untagged ORFeus with a GFP-AI cassette) was constructed by subcloning ORFeus under a Tet inducible promoter into a pCEP-puro vector^40^. pSS141 (pCEP ORFeus BSD-AI) and pSS207 (pCEP ORFeus-endo-dead BSD-AI) were constructed by replacing the GFP-AI cassette with BSD-AI. pSS131 (BSD-AI rescue construct) was constructed by replacing the L1PA1 in pBS-L1PA1CH_blast rescue (Addgene #69611) with tet-ORFeus and the backbone was replaced with pCEP-puro sequences.

### Primary siRNA screen

Quasi-stable cells expressing ORFeus GFP-AI were used for the screen. We plated 2,500 cells in each well of a 384-well plate; at the same time, cells were transfected with siRNA control or siRNA against specific proteins (Life Technologies). DharmaFECT transfection reagent (0.1 μL per well) (Dharmacon, prod. number T-2001-01) was used for siRNA transfection. 3 d after the knockdown, cells were fixed with Formalin (4% formaldehyde) and stained with DAPI for nuclear recognition. Plates were then scanned by the Arrayscan VTI and percentage of GFP positive cells was quantified by the built-in software.

Images were acquired (1 field per well, 2 channels per field, 5x magnification, 2×2 binning (3.7mm^2 field area)) on the ArrayScan VTI (Thermo Scientific) and analyzed simultaneously for percentage of GFP positive cells using the Target Activation Bioapplication (Thermo Scientific) Cellomics Scan version 6.6.0 (Build 8153) which identifies DAPI positive nuclei using dynamic isodata thresholding, minimal smoothing, and segmentation based on shape after background subtraction and quantifies the GFP intensity within the cytoplasm. Limits of fluorescence were set so that no cells were considered positive for preparations of cells not containing GFP.

A series of optimizations were conducted before implementing the method in a genome-wide setting: 1) optimization of Doxycycline induction time. L1 was induced for 3 d, at which point the retrotransposition rate reaches saturation (**Figure S1A**). 2) measurement and correlation of cell mortality and GFP fluorescence. To exclude the possibility of auto-fluorescence from dead cells, and to exclude the possibility that cells undergoing retrotransposition under these conditions had a higher rate of death, we labeled dead cells using a cell impermeant stain (NucRed, Life Technology, prod. number R37106). Notably, we found that the GFP and NucRed signals were mostly mutually exclusive, showing that the cells that underwent L1 retrotransposition were mostly alive at the time of measurement (**Figure S1B**). This observation minimized the possibility of false positive GFP measurements due to elevated mortality of the subset of cells that underwent L1 retrotransposition. Interestingly, in the subset of siRNAs that were used for optimization, we noticed increased cell death upon treatment with siRNAs that led to decreased retrotransposition (**Figure S1B**). This observation may indicate that siRNAs depleting essential genes induce cellular stress, that consequently (and perhaps indirectly), decreases L1 retrotransposition. As a practical solution for avoiding situations of this type, we set a minimum number of cells (1500 cells/well; the median number of cells/well across the entire dataset is 5228 cells/well) and eliminated all siRNAs that decreased the number of cells below this threshold. It is possible that L1 evolved to rely on essential cellular pathways (**Figure S1C**) and therefore siRNAs targeting essential genes are enriched in genes directly needed for L1 retrotransposition, in other words, host factors. These observations complicate the interpretation of results regarding those genes that decrease L1 retrotransposition.

The target activation (TA) image analysis algorithm uses DAPI to detect and identify a primary object (Nucleus) based on the difference in pixel intensities between the nucleus and the background (no nucleus). Once the primary object is identified the software draws a perimeter around the border of the nucleus creating a region of interest for the channel 1 object (ch1 ROI). The ch1 ROI is transferred into the GFP channel 2 (ch2 ROI) which is expanded to measure the average GFP expression in the cytoplasm. (Total GFP pixel Intensity in ch2 ROI / Total area of ch2 ROI). The diameter of the ch2 ROI is user defined to allow accurate modeling of the cytoplasm of each individual object (nuclei). In this screen we used ch1 ROI +8μm as ch2 ROI in order to capture the highest GFP signal.

### Data analysis of the whole-genome screen

We repeated each siRNA knockdown in triplicate in separate plates; control/mock siRNAs were included in each plate. A plate-based normalization was applied to each knockdown in order to minimize plate to plate differences. A source of plate variability also came from the number of passages of the reported cell line used, as the number of reporter plasmids slowly decreased or the reporter was silenced as cells were cultured. Cells within an empirically established window of passage numbers were therefore always used. The plate median was used to calculate the robust z-score of the log transformed %GFP=GFP/DAPI*100. To analyze the data, we first excluded wells showing low viability (< 1500 cells) as those data may bias results. We then defined inhibitors as those knockdowns having robust z-scores greater than 3 in at least two of the triplicates; supporters are defined as having robust z-scores smaller than −0.8 in at least two replicates or the percentage of GFP cells equals to 0 in all three replicates.

### Secondary validation screen

The settings of the 96 well validation followed the same procedures as the whole genome screen. We calculated the fold change of %GFP compared to siScramble for each knockdown.

### Exon-junction qPCR

Measuring L1 retrotransposition by genomic exon-junction qPCR analysis of spliced GFP we noticed that several measurements did not always agree with measurements of GFP expression. This discrepancy is particularly evident for the siRNAs that induced smaller effects on retrotransposition, which may underscore the differences between the two measurements; as previously reported a specific L1-GFP insertion may be quickly silenced after landing into heterochromatic locus^64^. These specific retrotranspositions cannot be detected with our imaging approach but can still be picked up by GFP-AI exon-junction qPCR, perhaps explaining some of the discrepancies. Contrary to these positive features of the qPCR measurement compared to GFP measurements, we observed that the evaluation of L1 retrotransposition by qPCR analysis appears to be a less sensitive assay, unable to identify factors with small effect on L1 activity. This could possibly be explained by the relatively low amount of spliced GFP DNA from cells that underwent retrotransposition among the rest of wild type genomic DNA.

After Arrayscan VTI analysis exon junction qPCR was performed, lysing the cells in the same wells by freezing and thawing in TE buffer. Cell lysates were then treated with 0.3μg/μl proteinase K (Invitrogen, prod. number 25530049) for 1 h at 37°C and ~50ng DNA was used for qPCR reaction. GFP and the control gene RPL21 were amplified using the following primers and probes:

GFP-F-JB17510: CAAGGTGAACTTCAAGATCC

GFP-R-JB17511: GCTCAGGTAGTGGTTGTC

RPL21-F-JB17589: AGCCTGCTCCACCCAGAGA

RPL21-R-JB17590: CCAGCAGCTCAGGCTCCTT

GFP probe: 6FAM-5’-TCGGCCAGCTGCAC-3’-BHQ1

RPL21 probe: HEX-CACACTTTGTGAGAACCA-BHQ1

Taqman probes were designed spanning the GFP exon junction (these anneal only to the intronless [i.e. post RNA-splicing] GFP region). The probe sequences were designed as previously described in^65^. The GFP expression was quantified by normalizing to the internal control, RPL21, using custom TaqMan FAM/HEX-labeled probes. Probes were synthesized by Integrated DNA Technologies (IDT).

We used LightCycler^®^ 480 Probes Master (Roche, prod. number 04707494001) for the qPCR reaction.

### CRISPR and BRCA1 rescue experiment

A gRNA (GGAGCCCACTTCATTAGTAC) targeting *BRCA1* was cloned into lentiCRISPR v2 (Addgene #52961) following Golden Gate cloning protocol^66^. Lentivirus carrying CRISPR-Cas9 constructs were produced using HEK293 cells with the following plasmids: pSS172 (pMD2.G, Addgene #12259), pSS173 (psPAX2, Addgene #12260), pSS164 (CRISPR Cas9 with BRCA1 gRNA). These three plasmids were co-transfected with Lipofectamine 2000 in 10cm plates. Viral supernatants were collected and filtered with 0.45μm filters. HCT116+ch3 cells (plated the previous day) were then infected with the collected lentivirus using 8μg/ml Hexadimethrine bromide. Infected cells were selected by 0.8μg/ml puromycin for > 3 d and used as a pool. Infection was optimized at and MOI of ~0.3. A *BRCA1* rescue construct (pSS146) was built by cloning the *BRCA1* cDNA into the ORFeus GFP-AI reporter plasmid (pEA79). Cells expressing integrated Cas9 and gRNA were than transfected with L1 reporter (with or without *BRCA1* cDNA) to measure the retrotransposition efficiency.

### Retrotransposition assay in 6 well plates

HeLa cells were plated into 6 well plates at 0.3 million cells per well. The next day, cells were transfected with L1 reporter construct using FuGENE HD Transfection Reagent (Promega, prod number E2311) following the standard protocol. 24 h after transfection, quasi-stable cells carrying the reporter plasmids were selected with 1μg/ml puromycin for > 5 d. Doxycycline (1μg/ml) was added to induce L1 expression. After 3 d, the percentage of GFP positive cells was measured by flow cytometry (FACS buffer: 1% FBS, 1mM EDTA and 100U/ml of Penicillin-Streptomycin) and dead cells were excluded from the analysis by 5μg/ml propidium iodide staining (ThermoFisher Scientific, prod. number P1304MP).

### Knockdown experiment and Aphidicolin treatment

HeLa cells were plated into 6 well plates with 0.3 million cells per well. The next day we transfected the cells with 7.5μl 10μM siRNA pools (3 siRNAs against same gene) using 100μl OPTIMEM (Gibco, prod. number 31985070) and 7.5μl Lipofectamine^®^ RNAiMAX Transfection Reagent (Life Technologies, prod. number 13778030). The next day, cells were replated into 6 well plates with 0.1 million cells per well. After 24 h, we added 1μg/ml doxycycline and various concentrations of Aphidicolin to the cells. After 24 h, we washed off Aphidicolin 3 times with PBS and cultured the cells in Doxycycline for another 2 d. After 2 d, we measured the retrotransposition efficiency by flow cytometry as described above. For a typical knockdown experiment, we add the doxycycline the next day after transfection and measure the retrotransposition by flow cytometry after 3 d.

### BSD-AI experiment

HeLa cells were first transfected with pSS141 (ORFeus BSD-AI) or pSS207 (endonuclease dead ORFeus BSD-AI) and quasi-stable cell lines were selected after 5 d of 1 μg/ml puromycin selection. We then performed knockdown with siRNA as described above. 24 h after knockdown, doxycycline was added for 3 d. We then plated 0.1 and 0.5 million cells for pSS141; 0.5 million and 5 million cells for pSS207 into 10cm plates under 15μg/ml Blasticidin selection. Cells were stained with crystal violet after 10 d for imaging.

### L1 insertion rescue

Quasi-stable cell line expressing pSS131 (ORFeus L1 rescue construct) was maintained under puromycin (1 μg/ml) selection for at least 5 d. We then performed knockdown experiment following the procedures described above. After 3 d of 1 μg/ml doxycycline treatment, cells were plated 1:10, 1:100 and 1:1000 in 10cm plates and kept under 15μg/ml Blasticidin selection. Single clones were hand-picked under microscope and then expanded. Each of the clones was expanded to reach confluency in a 6 cm plate. 3-5 million cells were then harvested and genomic DNA was isolated using QIAamp DNA mini kit (QIAGEN, prod. number 51304). The following L1 recovery steps were adapted from^24,52^ with modifications. 50μg of purified DNA was used for the following steps. 1) Genomic DNA was digested with 200U HindIII-HF (New England BioLabs, prod. number R3104S) at 37°C overnight. 2) The restriction enzyme was heat inactivated at 65C for 20min, followed by a self-ligation using 1200U T4 DNA ligase (New England BioLabs, prod. number M0202S) in 1ml total volume at room temperature overnight. 3) We then purified the DNA using Amicon Ultra 0.5ml, 50K (Millipore Sigma, prod number UFC505096) by centrifuging for 20 m at 14,000g and columns were washed with 500μl H2O twice. 4) We concentrated the DNA using ethanol precipitation into 5μl volume and pulled 5 samples into one transformation mix. 5) We then transformed the DNA into ElectroMAX™ DH1OB™ T1 R cells (Life Technologies, prod. number 12033-015) using electroporation and recovered the colonies on LB plates supplemented with 100μg/ml Blasticidin. 6) Each of the clones were then Sanger sequenced (Genewiz) using the following primers:

ss417 (detects 3’ junction): ATATATGAGTAACCTGAGGC

ss199 (detects 5’ junction): GGGCATCTTGAGCCCCTGCG

Additional primers (ss202-210) were used for longer insertions:

ss202: ATGGCCAAGCCTTTGTCTCA

ss203: GCTGAAGATGCGGTGCTTGG

ss204: GCTTCTTGGCGGCGTAGATG

ss205: GATCTCGCTGGGCTCGGTGC

ss206: GGGCGCTCACGATGGGGTTC

ss207: GTTCATCAGGCTGATGGGGC

ss208: TGCAGGGTCTTCTGGGTCTC

ss209: TGGTGCAGGGCGCTGTTCAG

ss210: GTTCTGCAGGGGCTGGTAGC

### Immunoblotting

HeLa M2 cells were lysed in SKL Triton lysis buffer (50 mM Hepes pH7.5, 150 mM NaCl, 1 mM EDTA, 1 mM EGTA, 10% glycerol, 1% Triton X-100, 25 mM NaF, 10 μM ZnCl2) supplemented with protease and phosphatase inhibitors (Complete-EDTA free, Roche/Sigma prod. number 11873580001; 1 mM PMSF and 1 mM NaVO4). NuPage 4XLDS sample buffer (ThermoFisher Scientific, prod. number NP0007) supplemented with 1.43M β-mercaptoethanol was added to the samples to reach a 1X dilution (350 mM β-mercaptoethanol final concentration) before gel electrophoresis performed using 4–12% Bis-Tris gels (ThermoFisher Scientific, prod. number WG1402BOX). Proteins were transferred on Immobilon-FL membrane (Millipore, prod. number IPFL00010), blocked for 1 hr with blocking buffer (LiCOR prod. number 927–40000): TBS buffer (50 mM Tris Base, 154 mM NaCl) 1:1 and then incubated with primary antibodies solubilized in LiCOR blocking buffer:TBS-Tween (0.1% Tween in TBS buffer) 1:1. Secondary goat antibodies conjugated to IRDye680 (anti-rabbit) or IRDye800 (anti-mouse) dyes (LiCOR prod. number 926–32210 and 926–68071), were used for detection of the specific bands on an Odyssey CLx scanner (LiCOR). 4H1 mouse monoclonal antibody targeting amino acids 35 to 44 of human ORF1p^40,67,68^ (Millipore, cat. number MABC1152) was used at 1:15000 dilution. ORF2-Flag were detected using FLAG-M2 antibody (Sigma, cat. number F1804) at 1:1000 dilution.

### Immunofluorescence

Immunofluorescence was performed as previously described^12^. We used the mouse *BRCA1* [D9] antibody (Santa Cruz cat. sc-6954) at 1:100 dilution and FLAG-M2 antibody (Sigma, cat. number F1804) at 1:1000 dilution for staining. Halotag7-ORF2p was detected using Halo tag ligand JF646^69^ as previously described^12^.

### IP-RT-qPCR and qPCR

HeLa-M2 cells expressing ORFeus (pLD401) from one 15 cm plate per IP were collected by trypsinization and lysed in SKL Triton lysis buffer (50 mM Hepes pH7.5, 150 mM NaCl, 1 mM EDTA, 1 mM EGTA, 10% glycerol, 1% Triton X-100, 25 mM NaF, 10 μM ZnCl2) supplemented with 1 mM DTT, 400 μM Ribonucleoside Vanadyl Complex (VRC) (NEB, prod. number S1402), 400U per ml of buffer of RNASEOUT (Thermo, prod. number 10777019), protein inhibitor and phosphatase inhibitor (Complete-EDTA free, Roche/Sigma prod. number 11873580001; 1 mM PMSF and 1 mM NaVO4). The cells-lysate was centrifuged for 15 m at 16,000 rcf at 4°C. Immunoprecipitations were conducted overnight at 4°C using mouse 4H1 ORF1p antibodies, mouse BRCA1 [D9] Santa Cruz cat. sc-6954) or control IgG from normal mouse (Santa Cruz sc-2025) coupled to magnetic beads (Dynabeads Antibody Coupling kit, Life Technologies, prod. number 14311D). Beads were washed five times in Triton buffer and resuspended in 100 μl of Triton buffer containing 30 μg of proteinase K (Invitrogen, prod. number 25530049). The mixture was incubated at 55°C for 30 min. 1 ml of Trizol (Life Technologies, prod. number 15596026) was directly added to the beads mixture and mRNA was purified using RNA clean-up and concentration columns (Norgen Biotek, prod. number 23600). cDNA was generated from RNA using the USB First-Strand cDNA synthesis kit for Real-Time PCR (Affimetrix, prod. number 75780). q-PCR was performed using a standard curve of pLD401 plasmid^40^. Each q-PCR reaction contained 2.5 μl of Sybr Green mastermix 2X (Roche, LightCycler 480 SYBR Green I Master, prod. number 04887352001), 25 nl of forward primer (100 μM), 25 nl of reverse primer (100 μM), 500 nl of cDNA for IP and water to 5 μl (final volume). qPCR was performed using a Light Cycler 480 (Roche) with standard conditions. The primers used for qPCR are reported below:

ORFeus ORF1:

JB13415 (forward): GCTGGATGGAGAACGACTTC

JB13416 (reverse): TTCAGCTCCATCAGCTCCTT

ORFeus ORF2:

JB13417 (forward): CTGATCAGCCGCATCTACAA

JB13418 (reverse): TGGTCTTGATCTGCATCTCG

BRCA1 (exon 5 and 6 spanning):

JB18182 (forward): TCAGCTTGACACAGGTTTGG

JB18183(reverse): GGATTTTCGGGTTCACTCTG RPL21:

JB17589 (forward): AGCCTGCTCCACCCAGAGA

JB17590 (reverse): CCAGCAGCTCAGGCTCCTT

### Live-cell imaging

Hela S.FUCCI cells expressing rtTA and stably expressing a pCEP-puro plasmid expressing ORFeus L1 with Halotag7-ORF2p and untagged ORF1p under a Tet inducible promoter^12^ were plated in 24 well plates at a density of 0.025-10^6 cells/well in 500ml of media. The next day 500μl of fresh media containing doxycycline 2μg/ml and Halo ligand JF646 200nM was added to the cells. After 4 h cells were moved to an EVOS-FL auto on stage incubator (Thermo-Fisher) for collection every 30 m for 48hr. Bright field was used for autofocus and GFP, RFP and cy5 cubes were used to collect fluorescence images of about 30 beacons per experiment.

## Supporting information

Movie1.FucciCell_ORF2_in_red_G1

Movie2.FucciCell_ORF2_in_green_S-G2

TableS1.screen_raw_data

TableS2.96well_validation

TableS3.L1_insertions

## DATA ACCESS

All the raw data of the primary and secondary screens are provided as supplemental tables.

## ACKNOWLEDGMENTS

We thank Tony Huang, Kathleen H. Burns for helpful discussions and comments on the manuscript. This work was supported by NIH grants P50GM107632 to J.D.B. and P01AG051449 to John Sedivy. The High Throughput Biology core is partially supported by Laura and Isaac Perlmutter Cancer Center Support Grant, “NIH/NCI P30CA16087” and NYSTEM Contract C026719.

## AUTHOR CONTRIBUTION

X.S., P.M. and J.D.B. conceived the project; X.S. and P.M., performed experiments; D.K. and C.Y. conducted the primary screen; D.L. and A.W. contributed new reagents/approaches; X.S., P.M., D.K., C.Y. and J.B. analyzed results; X.S., P.M. and J.D.B. wrote the manuscript; All authors read and commented on the manuscript.

---

**Figure S1.**
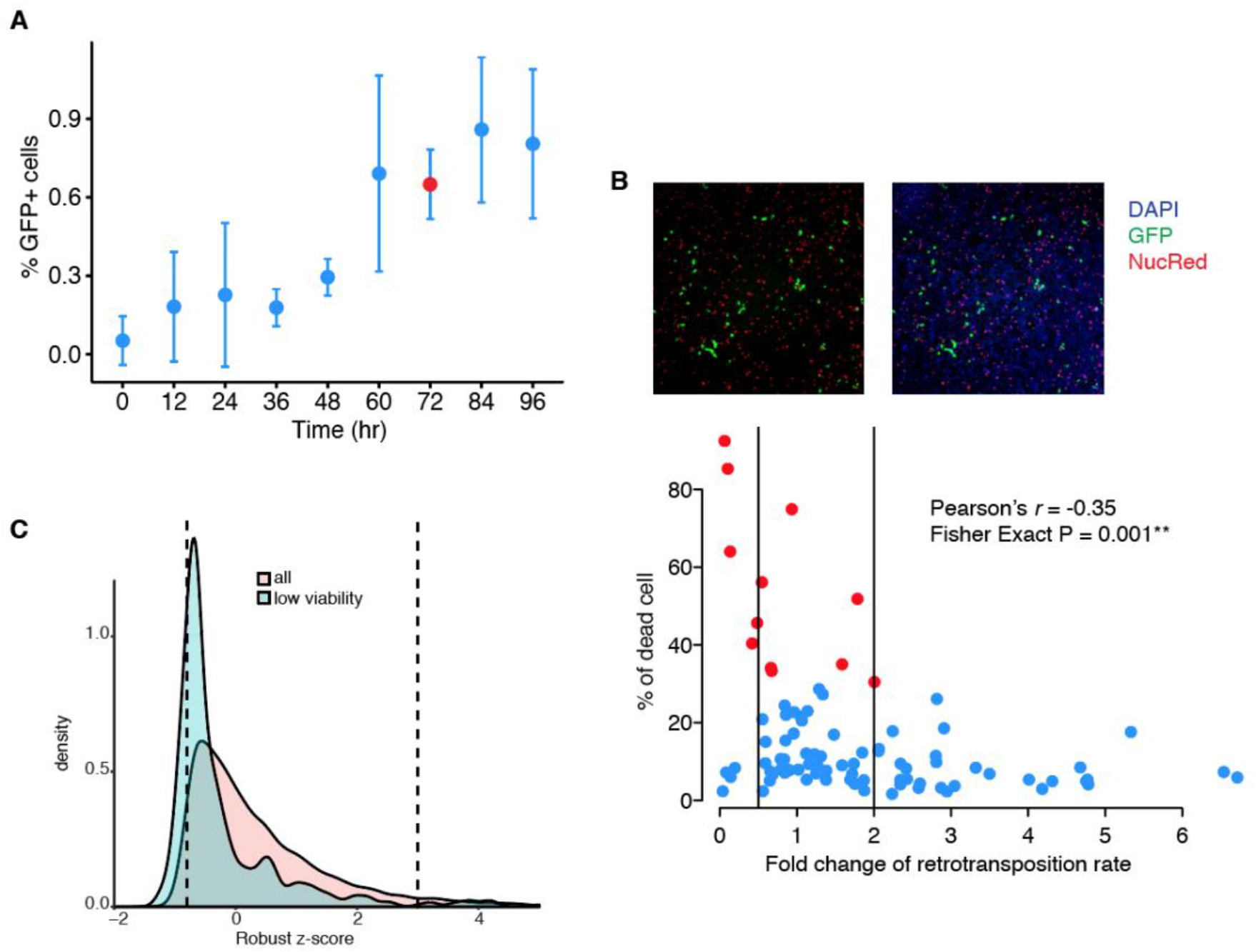
Optimization of the primary screen. **A.** Retrotransposition efficiency was measured over time by quantifying the percent of GFP positive cells. We used HeLa quasi-stable cells carrying ORFeus GFP-AI reporter and the L1 expression was induced by adding Doxycycline. Red dot marks the optimized time point (72 hours/3 d) utilized in the screen. Error bar: standard deviation of six biological replicates. **B.** We investigated the correlation between retrotransposition and cell death in a small set of knockdown samples. NucRed was used to stain dead cells and quantify the percentage of dead cells for each knockdown (upper panel). We found that death was negatively correlated with fold changes of retrotransposition efficiency (Pearson’s r= −0.35). We observed that high cell death occurred more often in cells that have decreased level of retrotransposition (Fisher Exact Test: p-value: 0.001). Red dots indicate samples with > 30% dead cells. Vertical bars indicate fold-change cutoffs for suppressors (foldchange <= 0.5) and enhancers (fold-change >=2) for retrotransposition. **C.** Distribution of the robust z-scores for siRNAs resulting in low cell viability (blue shade) compared to z-scores of the entire siRNA library (red shade). The dashed vertical lines indicate the thresholds set for inhibitors and supporters of L1 retrotransposition.

**Figure S2.**
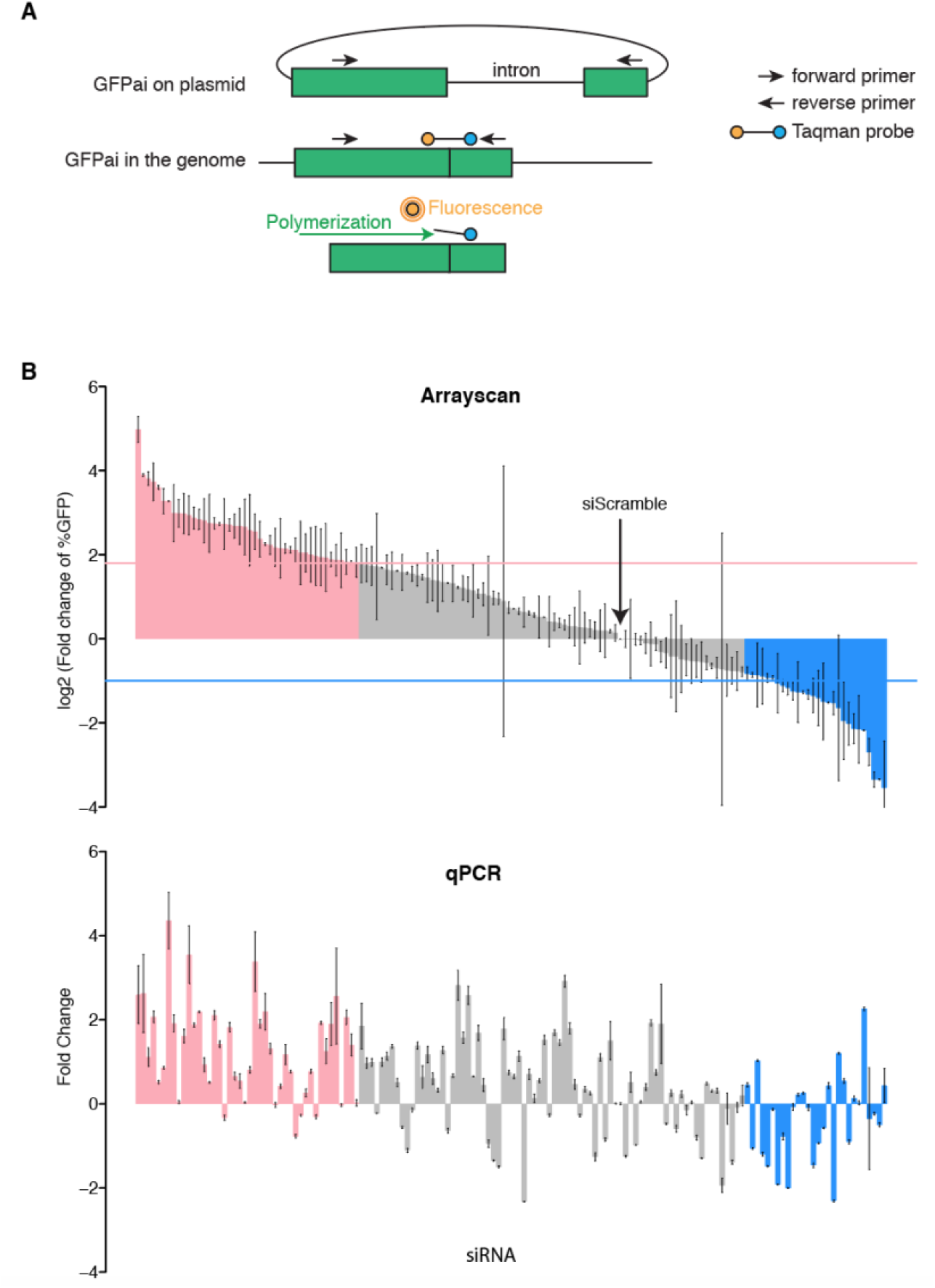
Junction qPCR method to evaluate L1 retrotransposition. **A.** Primers flanking the intron and a Taqman probe spanning the GFP-AI exon junctions were designed. Probes were labeled with a fluorophore and a quencher which inhibits fluorescence when both labels were in close proximity. During the qPCR reaction the exonuclease activity of the Taq polymerase degrades the annealed probes therefore releasing the fluorophore from the quencher and allowing measurement of fluorescence. **B.** Comparison of the 96 well validation assay and the junction qPCR assay. The same plates containing knockdown samples of the DNA repair factors analyzed by Arrayscan, were also used to measure L1 retrotransposition by junction qPCR. Fold changes of the qPCR quantification are plotted based on their Arrayscan measurement. Red horizontal line indicates the cutoff for the repressors (log2 fold change > 1.8). Blue horizontal line indicates the cutoff for the activators (log2 fold change < −1). Repressors are indicated by red bars and activators are indicated by blue bars. Error bars represent standard deviations from three biological replicates.

**Figure S3.**
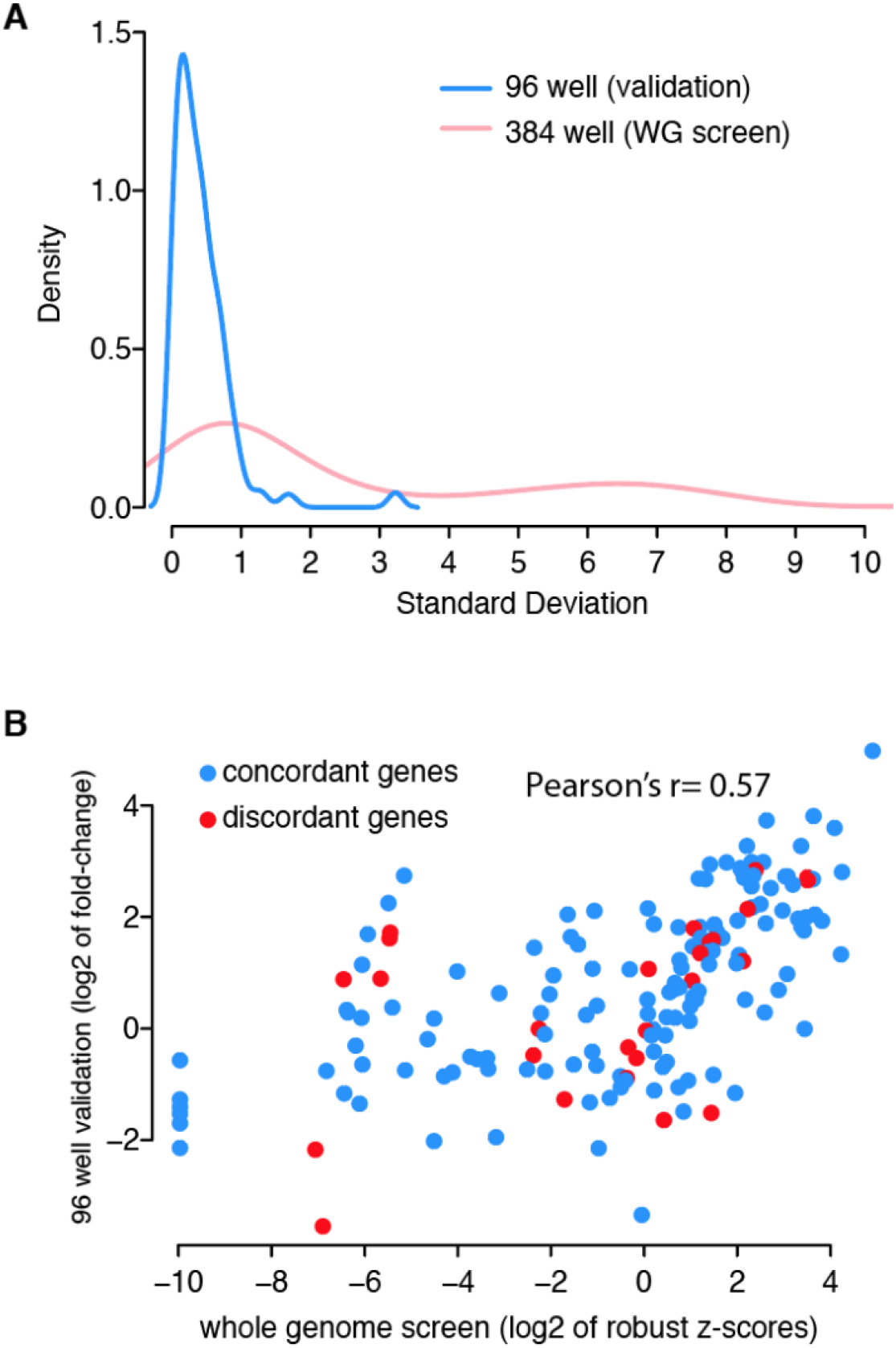
Comparison of the primary screen and 96 well validation. **A.** Distribution of the standard deviations from primary screen (384 well plates) and validation screen (96 well plates). **B.** Correlation of the fold changes from primary screen (384 well plates) and validation screen (96 well plates). Concordant genes and discordant genes were labeled as blue and red.

**Figure S4.**
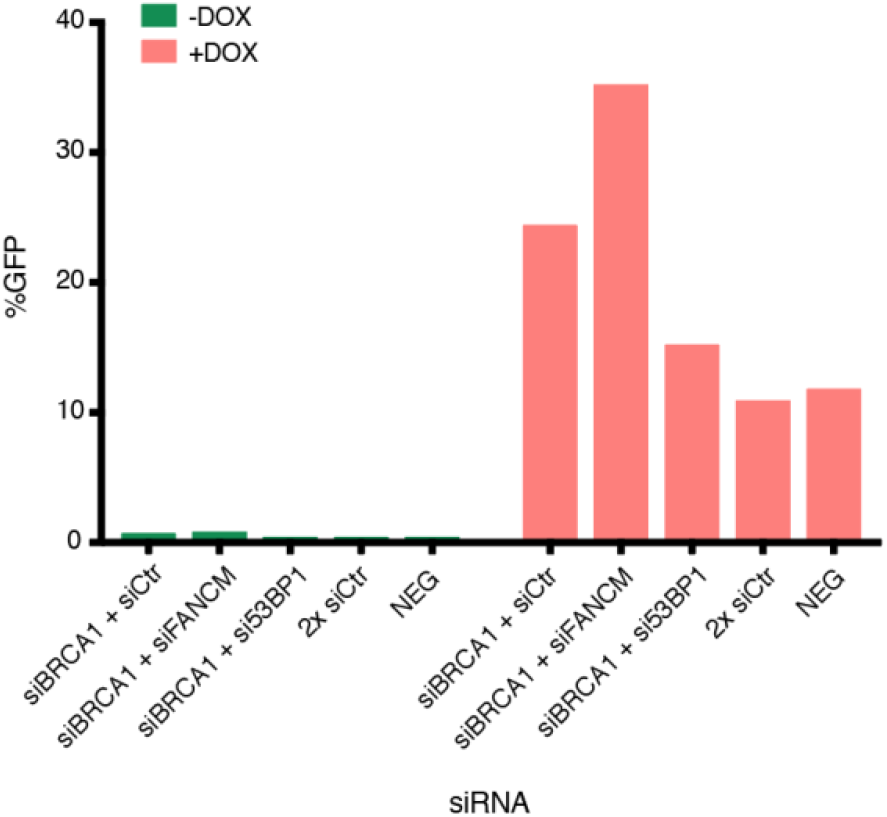
Measurements of L1RP retrotransposition in BRCA1 and 53BP1 depleted cells. Retrotransposition efficiency was measured as percentage of GFP positive cells using different combinations of siRNAs. GFP cells were counted with or without L1 induction (+/− Doxycycline).

**Figure S5.**
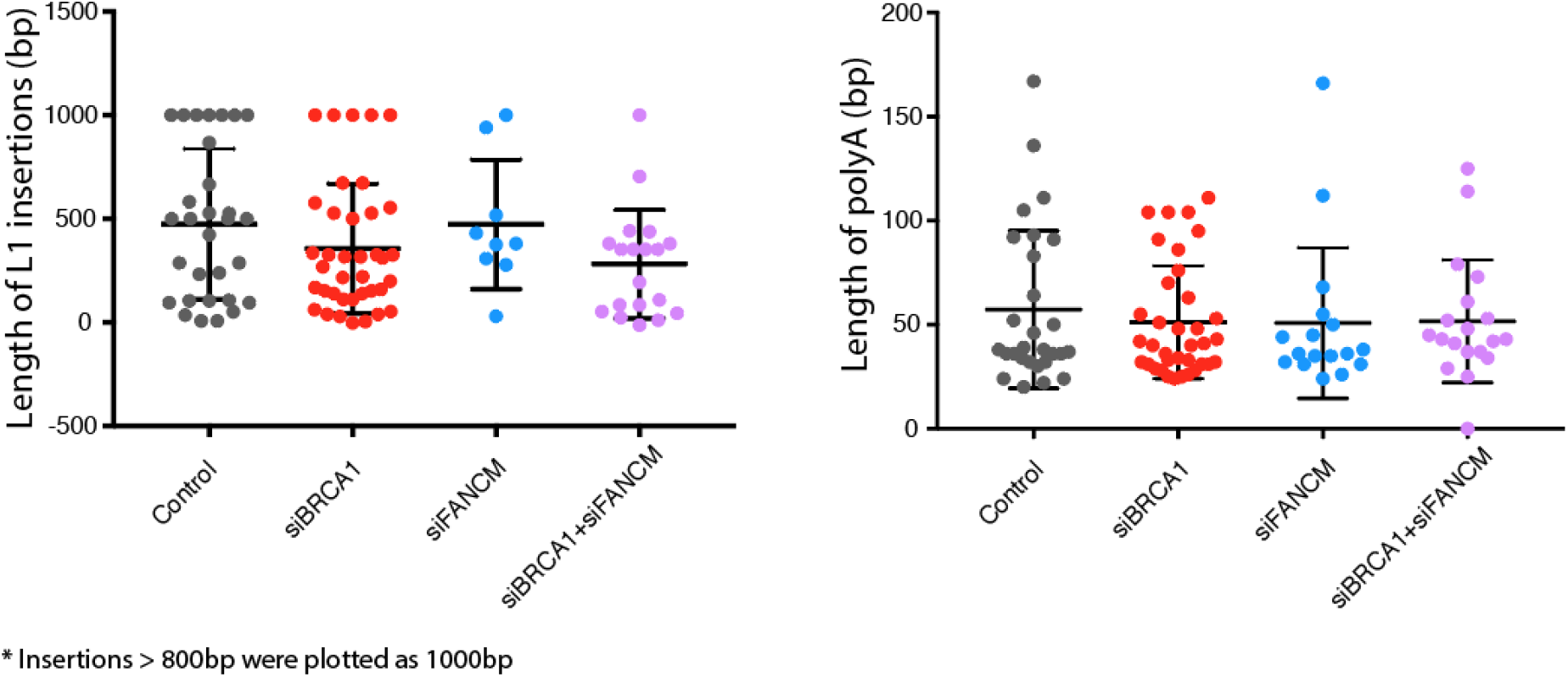
L1 insertion lengths and polyA lengths recovered in clones. Lengths of the sequenced L1 insertions were plotted in the left panel for control, BRCA1, FANCM and BRCA1-FANCM knockdown cells. Insertions larger than 800bp were plotted as 1000bp. Lengths of polyA from L1 insertions are plotted in the right panel.

**Figure S6.**
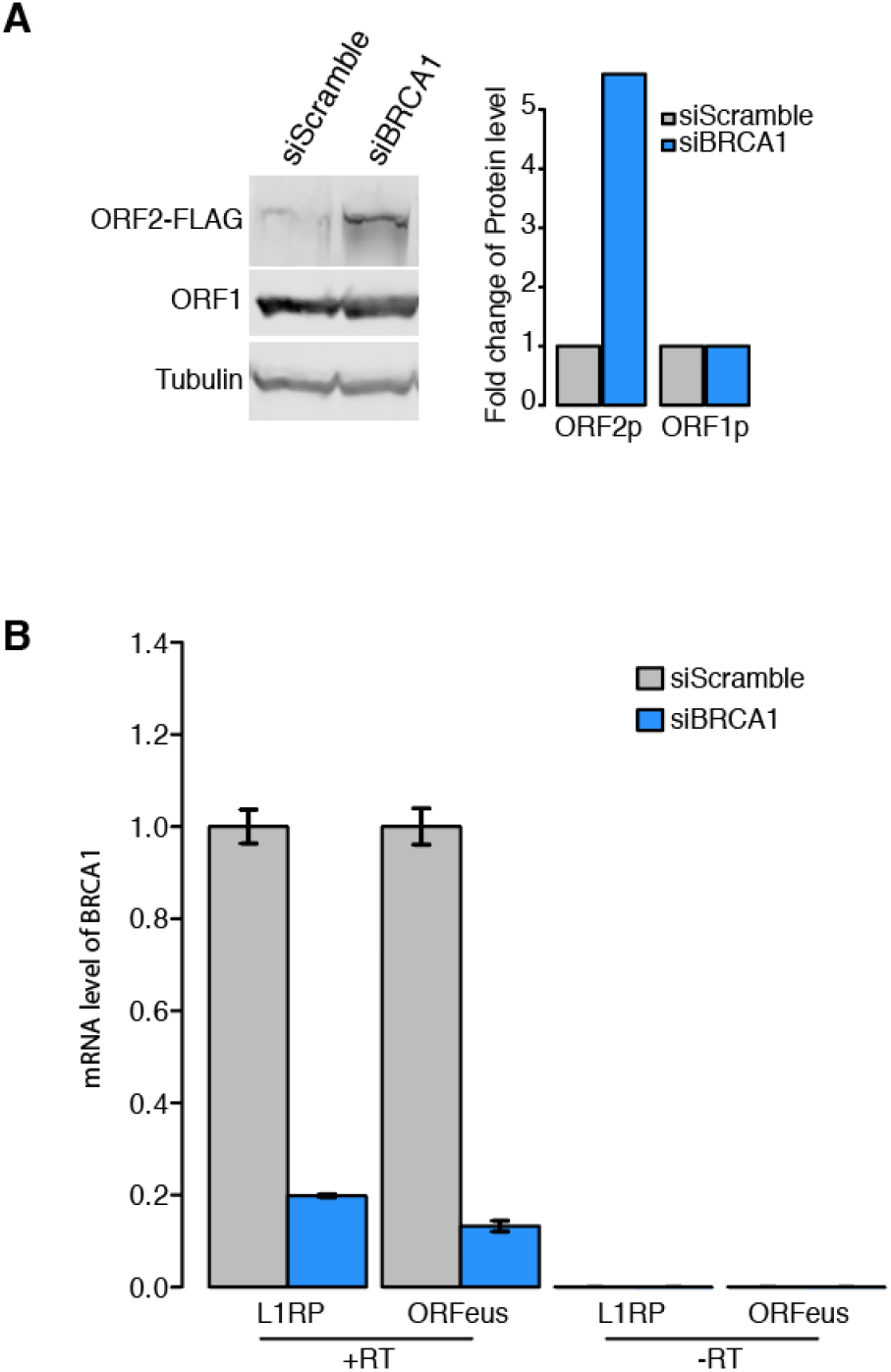
ORF2 protein level increases in BRCA1 knockdown cells. **A.** Western blot shows levels of ORF2 protein, ORF1 protein in control and BRCA1 knockdown cells expressing L1RP. To quantify the protein levels, we first normalized the signals to the level of Tubulin and then to the control cells (siScramble). **B.** Knockdown efficiency was measured by the abundance of BRCA1 mRNA in quasi-stable cells expressing either L1RP or ORFeus. We also ran the samples without reverse transcription to eliminate possible signal from genomic contaminations.

**Movie 1: Live-cell imaging of FUCCI cells expressing ORF2 in G1**

FUCCI cells expressing L1 were imaged every 30 minutes for 48 hours. Geminin and Cdt1 peptides are visualized in green and red respectively. Merged channels, ORF2p (cy5 channel) and bright field are shown as a movie. The merged channel (left panel) shows a cluster of cells located in the center of the field and starting to express ORF2p in G1 phase (red nuclei).

**Movie 2: Live-cell imaging of FUCCI cells expressing ORF2 in S/G2**

FUCCI cells expressing L1 were imaged every 30 minutes for 48 hours. Geminin and Cdt1 peptides are visualized in green and red respectively. Merged channels, ORF2p (cy5 channel) and bright field are shown as a movie. The merged channel (left panel) shows two cells in the center of the field and starting to express ORF2p in S/G2 phase (green nuclei).

**Table S1: Raw data of the genome-wide siRNA knockdown screen**

**Table S2: log2 fold changes of the DNA repairs factors in 96well validation screen**

**Table S3: L1 insertion sequences recovered in control and siBRCA1 cells**

## Reference

1. Burns, K. H. & Boeke, J. D. Human transposon tectonics. Cell 149, 740–752 (2012).

2. Huang, C. R. L., Burns, K. H. & Boeke, J. D. Active Transposition in Genomes. Annu. Rev. Genet. 46, 651–675 (2012).

3. Sassaman, D. M. et al. Many human L1 elements are capable of retrotransposition. Nat. Genet. 16, 37–43 (1997).

4. Brouha, B. et al. Hot L1s account for the bulk of retrotransposition in the human population. Proc. Natl. Acad. Sci. 100, 5280–5285 (2003).

5. Martin, S. L. & Bushman, F. D. Nucleic acid chaperone activity of the ORF1 protein from the mouse LINE-1 retrotransposon. Mol. Cell. Biol. 21, 467–475 (2001).

6. Cost, G. J., Feng, Q., Jacquier, A. & Boeke, J. D. Human L1 element target-primed reverse transcription in vitro. EMBO J. 21, 5899–5910 (2002).

7. Feng, Q., Moran, J. V, Kazazian, H. H. & Boeke, J. D. Human L1 Retrotransposon Encodes a Conserved Endonuclease Required for Retrotransposition. Cell 87, 905–916 (1996).

8. Kulpa, D. A. & Moran, J. V. Cis-preferential LINE-1 reverse transcriptase activity in ribonucleoprotein particles. Nat. Struct. Mol. Biol. 13, 655–660 (2006).

9. Wei, W. et al. Human L1 retrotransposition: cis preference versus trans complementation. Mol. Cell. Biol. 21, 1429–1439 (2001).

10. Alisch, R. S., Garcia-Perez, J. L., Muotri, A. R., Gage, F. H. & Moran, J. V. Unconventional translation of mammalian LINE-1 retrotransposons. Genes Dev. 20, 210–224 (2006).

11. Hohjoh, H. & Singer, M. F. Cytoplasmic ribonucleoprotein complexes containing human LINE-1 protein and RNA. EMBO J. 15, 630–639 (1996).

12. Mita, P. et al. LINE-1 protein localization and functional dynamics during the cell cycle. Elife 7, 210 (2018).

13. Luan, D. D., Korman, M. H., Jakubczak, J. L. & Eickbush, T. H. Reverse transcription of R2Bm RNA is primed by a nick at the chromosomal target site: A mechanism for non-LTR retrotransposition. Cell 72, 595–605 (1993).

14. Jurka, J. & Klonowski, P. Integration of Retroposable Elements in Mammals: Selection of Target Sites. J. Mol. Evol. (1996). doi:10.1007/BF02202117

15. Gilbert, N., Lutz, S., Morrish, T. A. & Moran, J. V. Multiple fates of L1 retrotransposition intermediates in cultured human cells. Mol. Cell. Biol. 25, 7780–7795 (2005).

16. Moran, J. V et al. High frequency retrotransposition in cultured mammalian cells. Cell 87, 917–927 (1996).

17. Symer, D. E. et al. Human l1 retrotransposition is associated with genetic instability in vivo. Cell 110, 327–338 (2002).

18. Ostertag, E. M. & Kazazian Jr, H. H. Biology of Mammalian L1 Retrotransposons. dx.doi.org 35, 501–538 (2003).

19. Szak, S. T., Pickeral, O. K., Makalowski, W. & Boguski, M. S. Molecular archeology of L1 insertions in the human genome. Genome … (2002).

20. Morrish, T. A. et al. DNA repair mediated by endonuclease-independent LINE-1 retrotransposition. Nat. Genet. 31, 159–165 (2002).

21. Morrish, T. A. et al. Endonuclease-independent LINE-1 retrotransposition at mammalian telomeres. Nature 446, 208–212 (2007).

22. Sen, S. K., Huang, C. T., Han, K. & Batzer, M. A. Endonuclease-independent insertion provides an alternative pathway for L1 retrotransposition in the human genome. Nucleic Acids Res. 35, 3741–3751 (2007).

23. Burma, S., Chen, B. P. C. & Chen, D. J. Role of non-homologous end joining (NHEJ) in maintaining genomic integrity. DNA Repair (Amst). 5, 1042–1048 (2006).

24. Servant, G. et al. The Nucleotide Excision Repair Pathway Limits L1 Retrotransposition. Genetics 205, 139–153 (2017).

25. Han, K. et al. Genomic rearrangements by LINE-1 insertion-mediated deletion in the human and chimpanzee lineages. Nucleic Acids Res. 33, 4040–4052 (2005).

26. Rodriguez-Martin, B. et al. Pan-cancer analysis of whole genomes reveals driver rearrangements promoted by LINE-1 retrotransposition in human tumours. bioRxiv 179705 (2017). doi:10.1101/179705

27. Hancks, D. C. & Kazazian, H. H. Active human retrotransposons: variation and disease. Curr. Opin. Genet. Dev. 22, 191–203 (2012).

28. Belgnaoui, S. M., Gosden, R. G., Semmes, O. J. & Haoudi, A. Human LINE-1 retrotransposon induces DNA damage and apoptosis in cancer cells. Cancer Cell Int. 6, 13 (2006).

29. Farkash, E. A. & Luning Prak, E. T. DNA damage and L1 retrotransposition. J. Biomed. Biotechnol. 2006, 37285–37288 (2006).

30. Farkash, E. A., Kao, G. D., Horman, S. R. & Prak, E. T. L. Gamma radiation increases endonuclease-dependent L1 retrotransposition in a cultured cell assay. Nucleic Acids Res. 34, 1196–1204 (2006).

31. Gasior, S. L., Wakeman, T. P., Xu, B. & Deininger, P. L. The Human LINE-1 Retrotransposon Creates DNA Double-strand Breaks. J. Mol. Biol. 357, 1383–1393 (2006).

32. An, W. L1 expression and regulation in humans and rodents. Front. Biosci. E4, 2203–2225 (2012).

33. Knijnenburg, T. A. et al. Genomic and Molecular Landscape of DNA Damage Repair Deficiency across The Cancer Genome Atlas. Cell Rep. 23, 239–254.e6 (2018).

34. Martin, S. A., Lord, C. J. & Ashworth, A. DNA repair deficiency as a therapeutic target in cancer. Curr. Opin. Genet. Dev. 18, 80–86 (2008).

35. Coufal, N. G. et al. Ataxia telangiectasia mutated (ATM) modulates long interspersed element-1 (L1) retrotransposition in human neural stem cells. Proc. Natl. Acad. Sci. U. S. A. 108, 20382–20387 (2011).

36. Suzuki, J. et al. Genetic Evidence That the Non-Homologous End-Joining Repair Pathway Is Involved in LINE Retrotransposition. PLoS Genet. 5, e1000461 (2009).

37. Brégnard, C. et al. Upregulated LINE-1 Activity in the Fanconi Anemia Cancer Susceptibility Syndrome Leads to Spontaneous Pro-inflammatory Cytokine Production. EBioMedicine 8, 184–194 (2016).

38. Liu, N. et al. Selective silencing of euchromatic L1s revealed by genome-wide screens for L1 regulators. Nature (2017). doi:10.1038/nature25179

39. Hampf, M. & Gossen, M. Promoter Crosstalk Effects on Gene Expression. J. Mol. Biol. (2007). doi:10.1016/j.jmb.2006.10.009

40. Taylor, M. S. et al. Affinity Proteomics Reveals Human Host Factors Implicated in Discrete Stages of LINE-1 Retrotransposition. Cell 155, 1034–1048 (2013).

41. An, W. et al. Characterization of a synthetic human LINE-1 retrotransposon ORFeus – Hs. Mob. DNA 2, 1 (2011).

42. Krejci, L., Altmannova, V., Spirek, M. & Zhao, X. Homologous recombination and its regulation. Nucleic Acids Research (2012). doi:10.1093/nar/gks270

43. Li, X. & Heyer, W.-D. Homologous recombination in DNA repair and DNA damage tolerance. Cell Res. (2008). doi:10.1038/cr.2008.1

44. Bunting, S. F. et al. 53BP1 inhibits homologous recombination in Brca1-deficient cells by blocking resection of DNA breaks. Cell 141, 243–254 (2010).

45. Polato, F. et al. CtIP-mediated resection is essential for viability and can operate independently of BRCA1. J. Exp. Med. 211, 1027–1036 (2014).

46. Ceccaldi, R., Sarangi, P. & D’Andrea, A. D. The Fanconi anaemia pathway: new players and new functions. Nat. Rev. Mol. Cell … 17, 337–349 (2016).

47. Michl, J., Zimmer, J. & Tarsounas, M. Interplay between Fanconi anemia and homologous recombination pathways in genome integrity. EMBO J. 35, 909–923 (2016).

48. Ruffner, H., Joazeiro, C. A., Hemmati, D., Hunter, T. & Verma, I. M. Cancer-predisposing mutations within the RING domain of BRCA1: loss of ubiquitin protein ligase activity and protection from radiation hypersensitivity. Proc. Natl. Acad. Sci. 98, 5134–5139 (2001).

49. Shen, S. X. et al. A targeted disruption of the murine Brca1 gene causes gamma-irradiation hypersensitivity and genetic instability. Oncogene 17, 3115–3124 (1998).

50. Fungtammasan, A., Walsh, E., Chiaromonte, F., Eckert, K. A. & Makova, K. D. A genome-wide analysis of common fragile sites: What features determine chromosomal instability in the human genome? Genome Res. 22, 993–1005 (2012).

51. Arlt, M. F. et al. BRCA1 is required for common-fragile-site stability via its G2/M checkpoint function. Mol. Cell. Biol. 24, 6701–6709 (2004).

52. Gilbert, N., Lutz-Prigge, S. & Moran, J. V. Genomic deletions created upon LINE-1 retrotransposition. Cell 110, 315–325 (2002).

53. Jensen, S., Gassama, M. P. & Heidmann, T. Retransposition of the Drosophilia LINE I Element Can Induce Deletion in the Target DNA: A Simple Model Also Accounting for the Variability of the Normally Observed Target Site Duplications. Biochem. Biophys. Res. Commun. 202, 111–119 (1994).

54. Dacheux, E. et al. BRCA1-Dependent Translational Regulation in Breast Cancer Cells. PLoS One 8, e67313 (2013).

55. Robbez-Masson, L. et al. The HUSH complex cooperates with TRIM28 to repress young retrotransposons and new genes. Genome Res. 28, 836–845 (2018).

56. Mao, Z., Bozzella, M., Seluanov, A. & Gorbunova, V. DNA repair by nonhomologous end joining and homologous recombination during cell cycle in human cells. Cell Cycle 7, 2902–2906 (2014).

57. Rothkamm, K., Kruger, I., Thompson, L. H. & Lobrich, M. Pathways of DNA Double-Strand Break Repair during the Mammalian Cell Cycle. Mol. Cell. Biol. 23, 5706–5715 (2003).

58. Symington, L. S. Mechanism and regulation of DNA end resection in eukaryotes. Crit. Rev. Biochem. Mol. Biol. 51, 195–212 (2016).

59. Mita, P. & Boeke, J. D. Cycling to Maintain and Improve Fitness: Line-1 Modes of Nuclear Entrance and Retrotransposition. SLAS Discov. (2018). doi:10.1177/2472555218767842

60. Jacobs, J. Z. et al. Arrested replication forks guide retrotransposon integration. Science (80-.). (2015). doi:10.1126/science.aaa3810

61. Tubbs, A. et al. Dual Roles of Poly(dA:dT) Tracts in Replication Initiation and Fork Collapse. Cell (2018). doi:10.1016/j.cell.2018.07.011

62. Moldovan, G.-L. & D’Andrea, A. D. How the Fanconi Anemia Pathway Guards the Genome. Annu. Rev. Genet. (2009). doi:10.1146/annurev-genet-102108-134222

63. Chaudhuri, A. R. et al. Replication fork stability confers chemoresistance in BRCA-deficient cells. Nature (2016). doi:10.1038/nature18325

64. Garcia-Perez, J. L. et al. Epigenetic silencing of engineered L1 retrotransposition events in human embryonic carcinoma cells. Nature 466, 769–773 (2010).

65. Xie, Y. et al. Cell division promotes efficient retrotransposition in a stable L1 reporter cell line. Mob. DNA 4, 10 (2013).

66. Sanjana, N. E., Shalem, O. & Zhang, F. Improved vectors and genome-wide libraries for CRISPR screening. Nat. Methods 11, 783–784 (2014).

67. Rodić, N. et al. Long interspersed element-1 protein expression is a hallmark of many human cancers. Am. J. Pathol. 184, 1280–1286 (2014).

68. Doucet-O’Hare, T. T. et al. LINE-1 expression and retrotransposition in Barrett’s esophagus and esophageal carcinoma. Proc. Natl. Acad. Sci. (2015). doi:10.1073/pnas.1502474112

69. Grimm, J. B. et al. A general method to improve fluorophores for live-cell and singlemolecule microscopy. Nat. Methods (2015). doi:10.1038/nmeth.3256

