## Supplementary material for "BRCA1 Mediated Homologous Recombination and S Phase DNA Repair Pathways Restrict LINE-1 Retrotransposition in Human Cells": TableS3.L1_insertions

*Insertion size was counted upstream of BSD

*Insertion strand is which genome strand L1 inserts

*Target site duplication

*Target site deletion

siScramble:

Clone C1 - TSD

- Cleavage site: CAAA/AACT
- Insertion site: chr7: 89405027
- Insertion strand: F
- polyA length: 39bp
- Insertion size: 107bp
- GAAAAGTTTCTGAGGCAAAAACTTATAACCTGA-L1-pA-AACTTATAACCTGAAGGTTTAATAGGGTAAATGAAAAA

Clone C2 - TSD

- Cleavage site: TTTT/AAAA
- Insertion site: chr2: 214995030
- Insertion strand: R
- polyA length: 34bp
- Insertion size: 867bp
- GTTTTGTTCTGCACTTTGCTTTTAAAAATT-L1-pA-AAAAATTAAAATACTGAAGCTATAAAAATCTG

Clone C3 – No TSD

- Cleavage site: CT/AAGAGC
- Insertion site: chr22: 16599808
- Insertion strand: R
- polyA length: 50bp
- Insertion size: 35bp
- CTCCTGCTGGAGAGTTCACAAGAAACAAAGGGTACTAA-L1-pA-GAGCAAAAAAGTGAGCC

Clone C4 - TSD

- Cleavage site: TCAA/AATG
- Insertion site: chr2: 27854824
- Insertion strand: R
- polyA length: 36bp
- Insertion size: 500bp
- GATGTATGCATGTATTCAAACCCATCAAAATGTATA-L1-pA-AATGTATACTTTAAATATATGCAGGGTTTTTAT

Clone C5 - TSD

- Cleavage site: TCAA/AATG
- Insertion site: chr12: 39216051
- Insertion strand: F
- polyA length: 22bp
- Insertion size: 51bp
- GCAACATTAAACAATAGAAGA-L1-pA-AACAATAGAAGACAGCAGG

Clone C6 – TSD

- Cleavage site: TCAA/AATG
- Insertion site: chrX: 18769352
- Insertion strand: R
- polyA length: 111bp
- Insertion size: 666bp
- ATTTCAAACTGGTCACTCTAATTAAGCAAC-L1-pA-AACCCAAAAAAAGCCC

Clone C7 - TSD

- Cleavage site: AAAAAA/TT
- Insertion site: chr15: 75907956
- Insertion strand: R
- polyA length: 46bp
- Insertion size: 233bp
- GGTGGACTTTAAAAAAATTTTTAAAATTTT-L1-pA-AAAAAAATTTTTTTTTTTTGAGACACAGTCTCACTCTTTAT

Clone C8 – TSD with insertion

- Cleavage site: AAAG/AGAT
- Insertion site: chr1: 30562234
- Insertion strand: F
- polyA length: 36bp
- Insertion size: 7bp
- GGCAAATTACATCAAGCAAAATTCAAAAGtctttt-L1-pA-AAAAGAGATCATTAATTGAACTTA

Clone C9 - Inversion

- Cleavage site: TTTT/AGAA
- Insertion site: chr10: 2188834
- Insertion strand: R
- polyA length: 83bp
- Insertion size: 287bp
- TTTGGAAATCATTCACCATTTTGTATAGTATTTTAGAAA-L1(with inversion)-pA-AGAAATTTAATTGTTTGGAAATCATTCACC

Clone C10 – TSD

- Cleavage site: TTTT/AGAA
- Insertion site: chr17: 58336051
- Insertion strand: R
- polyA length: 37bp
- Insertion size: 311bp
- CAAAGAGCCAACAAGAGAGAAGCCACA-L1-pA-GAGAGAAGCCACAGACTTAACCACT

Clone C11 – Inversion

- Cleavage site: AAA/ACCAA
- Insertion site: chr3: 148488677
- Insertion strand: F
- polyA length: 30bp
- Insertion size: 582bp
- CAGAGCGAGACTCCGTCTCAAAAAAAAAAAAAAAAAAAAAAAAAAAA-L1(with inversion)-pA-AACCAAAATACTTGTTATAACTATAAAATGTTTTGAGTAAG

Clone C12 - Inversion

- Cleavage site: AAAA/AGAA
- Insertion site: chr13: 51225976
- Insertion strand: R
- polyA length: 20bp
- Insertion size: 239bp
- AAACAAGTTAACAACAAAAAAAAAAAAAAAAAAA-L1(with inversion)-pA-AAAAGAAAGCAAGCAAGCAACATTACAGATTATCATCTTTTC

Clone C13 – Inversion (No TSD)

- Cleavage site: GCAT/GAAA
- Insertion site: chr1: 205590392
- Insertion strand: F
- polyA length: 55bp
- Insertion size: 317bp
- GACTTTCATATTTTTACTGCAT-L1(with Inversion)-pA-GAAAGAAGTCTTTTGTCTTTCCATTTCCAT

Clone C14 - inversion

- Cleavage site: AGTT/CAAA
- Insertion site: chr13: 25122804
- Insertion strand: F
- polyA length: 36bp
- Insertion size: 501bp
- CATAATGGTAACTACCACAATGTGTTGAGAATTAGGCAAAATA-L1(with inversion)-pA-CAAAATAACTTGTTATAACTATAAAATGTTTTGAGTAA

Clone C15 – inversion (No TSD)

- Cleavage site: CATT/TAAA
- Insertion site: chr2: 182044868
- Insertion strand: R
- polyA length: 24bp
- Insertion size: 527bp
- ATGGTGATAAAAGAGAAAAATAGAATAAAAACACATT-L1-pA-TAAATGCAGTTTGTAGATTAACGCCTATGATAATCTGTGTCTTATATGTA

Clone C16 - TSD

- Cleavage site: TTTT/AAAA
- Insertion site: chr13: 71447522
- Insertion strand: F
- polyA length: 37bp
- Insertion size: 423bp
- TAGAGTATTGAACAGATTGATCATTTATTTTAAAA-L1-pA-AAAACCTAATGAAGTTTGTTNAANGCCAATCATATGGCCGGGCGCAGTGGCT

Clone C17 - TSD

- Cleavage site: CATT/AAAT
- Insertion site: chr2: 182044850
- Insertion strand: R
- polyA length: 24bp
- Insertion size: 527bp
- CTCATAATGGTGATAAAAGAGAAAAATAGAATAAAAACACATTAAAT-L1-pA-TAAATGCAGTTTGTAGATTAACGCCTATGATAA

siBRCA1:

Clone B1 - Inversion

- Cleavage site: CATT/TAAA
- Insertion site: chr2: 182044868
- Insertion strand: R
- polyA length: 24bp
- Insertion size: 527bp
- AGAATAAAAACACATTTAAATGCAGTTTG-L1(with inversion)-pA-TAAATGCAGTTTGTAGATTAACGCCTATGATAATCTGTGT

Clone B2 – Inversion (no TSD)

- Cleavage site: TAAC/CTTC
- Insertion site: chr11: 24288586
- Insertion strand: F
- polyA length: 33bp
- Insertion size: 326bp
- CAAGTTTTCTTTGTTACATTTTAATAGTTAAC-L1(with inversion)-pA-CTTCCTGTATATTGAAACAATACA

Clone B3 – Target site deletion

- Cleavage site: AACA/GAAC
- Insertion site: chr2: 233523673
- Insertion strand: R
- polyA length: 76bp
- Insertion size: 139bp
- ATAGATTTCTCCTAAAAAAATAAAGATAA-L1-pA-GAAAGTTTCAAAAGAAAAATC

Clone B4 - Inversion

- Cleavage site: TT/AAGAAA
- Insertion site: chr4: 8438589
- Insertion strand: F
- polyA length: 43bp
- Insertion size: 268bp
- CTAAAGGAATTAACTTTAAGAAAGCAAAT-L1(with inversion)-pA-GAAAGCAAATGCAGGC

Clone B5 - TSD

- Cleavage site: CTT/AGAAG
- Insertion site: chr12: 7261302
- Insertion strand: R
- polyA length: 31bp
- Insertion size: 153bp
- AACAATCATTTCCTTAGAAGCGAG-L1-pA-GAGGCGAGCCTTGCAGTT

Clone B6 - Target site deletion

- Cleavage site: GCTG/TTAA
- Insertion site: chr13: 51225976
- Insertion strand: R
- polyA length: 20bp
- Insertion size: 239bp
- GGGTGACAGAGAGACCTGTCTGCTGCTGCTGCTGTTAAAAAAAAAAAAAAAAAAAAAAAAAAAAAAAAAAA-L1-pA-GAAAGCAAGCAAGCAACATTACAGATTATCATCTTT

Clone B7 – Target site deletion

- Cleavage site: CCAA/TAAT
- Insertion site: chrX: 80758275
- Insertion strand: F
- polyA length: 38bp
- Insertion size: 501bp
- ATCGAGGCCATCCTGGCTAACACGGTGAAACCCCGTCTCTACTAAAAATACAAAAAATTAGCCGGGTGTGGTAGCGGGCGCCTGTAGTCCCAGCTACTCGGGAGGCTGAGGCAGGAGAATGGCGTGAACCCGGGAGGCGGAGCTTGCAGTGAGCCGAGATCGCGCCACTGCACTCCAGCCTGGGCGACAGAGCGAGACTCCGTCTCAAAAAAAAAAAAAAAAAAAAAAAAAAAA-L1-pA-GAACTTGAAAATAATATCTATTCCATTTGGC

Clone B8 - Target site deletion

- Cleavage site: ACTA/TAAT
- Insertion site: chr6: 151709712
- Insertion strand: R
- polyA length: 32bp
- Insertion size: 577bp
- ATAGAAAAAATGCTCAGCATCACTATAATCATCA-L1pA-GAAAATGGAAATTAAAACCACAAAGAG

Clone B9 - Target site deletion

- Cleavage site: CTAC/CCAG
- Insertion site: chrX: 53034290
- Insertion strand: F
- polyA length: 34bp
- Insertion size: 311bp
- TTTTGAAAAAGAACAAAGTTAGAGACTTACTCTACCCAGTTTTTATCTACAGTAATCAACATGGTGAGATGTTGGTGTAAAGATGAATGAAAAAGAAGAGAGTCAA-L1-pA-GAAATAGGCCTACACATATAT

Clone B10 - Target site deletion

- Cleavage site: ACAG/ATGG
- Insertion site: chr11: 124358106
- Insertion strand: F
- polyA length: 36bp
- Insertion size: 39bp
- GATAAAGAAGTGGGTAACAGATGGCCACACAGCAATCATAGGCCATTGACGTCAGCATATAACATTCAGAAATAAATTAAAAAAAAAAAAAAA-L1-pA-CAGAAAAAGAAAAAGAAAAGCTAGAT

Clone B11 - TSD

- Cleavage site: AAAA/ACTC
- Insertion site: chr7: 106486541
- Insertion strand: F
- polyA length: 26bp
- Insertion size: 5bp
- TTCATTTCAATGATTAATCAAGAAAAAACTC-L1-pA-CTCAATGATTAATCATTTCAATAC

Clone B12 – Inversion (No TSD)

- Cleavage site: GTAC/TGGA
- Insertion site: chr11: 8500184
- Insertion strand: F
- polyA length: 48bp
- Insertion size: 199bp
- ATATCTACTTTCACCATTTCTATTTAAGATTGTAC-L1-pA-TGGAGGTCCTACACACTACACAGCACCTGGAGGTCCTACNCTACAGCACAATT

Clone B13 - Inversion

- Cleavage site: CACA/TTAA
- Insertion site: chr2: 182044852
- Insertion strand: R
- polyA length: 40bp
- Insertion size: 527bp
- GATAAAAGAGAAAAATAGAATAAAAACACATT-L1(with inversion)-pA-TTAANTGCNGTTTGTAAATTAACGCCTATGATAATCTG

Clone B14 – TSD (super long)

- Cleavage site: TAAA/AGTG
- Insertion site: chr17: 42272150
- Insertion strand: F
- polyA length: 63bp
- Insertion size: -2bp
- ACATTTATATTAAAagtgacctatcacttaagtgatccaggagagtaaaggaagagttttaaggctgaaggtgattcctcctgagctctaataccataccctgaatttcaccttgttcttaactcttttccatgacactgtgtatggcttttaagcgagtccagaaaaacggctatcattactacttaatataaaccatggggttctgtatcatgtaaacccagagcaactcctctttgaagcaccgacattcaag-L1-pA-agtgacctatcacttaagtgatccaggagagtaaaggaagagttttaaggctgaaggtgattcctcctgagctctaataccataccctgaatttcaccttgttcttaactcttttccatgacactgtgtatggcttttaagcgagtccagaaaaacggctatcattactacttaatataaaccatggggttctgtatcatgtaaacccagagcaactcctctttgaagcaccgacattcaagGCTCTACATAACATGTTT

Clone B15 – No TSD

- Cleavage site: AGTG/GGGG
- Insertion site: chr17: 42272152
- Insertion strand: F
- polyA length: 29bp
- Insertion size: 111bp
- TCCTGCAACATTTATATTAAAAGTG-L1-pA-GGGGCCCTTCCCTT

Clone B16 - inversion

- Cleavage site: TAAC/ACTT
- Insertion site: chr11: 24288586
- Insertion strand: F
- polyA length: 42bp
- Insertion size: 326bp
- TTTCTTTGTTACATTTTAATAGTTAAC-L1(with inversion)-pA-ACTTCCTGTATATTGAAACAATACA

Clone B17 – No TSD

- Cleavage site: GCG/AGCCT
- Insertion site: chr12: 7261335
- Insertion strand: R
- polyA length: 31bp
- Insertion size: 53bp
- TATATTTACAAAACAATCATTTCCTTAGAAGCG-L1-pA-gaggcgAGCCTTGCAGTTTCTATTTTCTCATGCCCCTTTACATAGT

Clone B18 - inversion

- Cleavage site:
- Insertion site: chr5: 101249190
- Insertion strand: F
- polyA length: 41bp
- Insertion size: 62bp
- ATAGAGACATAACTACTAGCTTTATA-L1(with inversion)-pA-GAAAGAGGAAAGAATGTTAANANAATGCTAAAAACAATT

Clone B19 – Target site deletion

- Cleavage site: AATA/AAAG
- Insertion site: chr13: 25122782
- Insertion strand: F
- polyA length: 38bp
- Insertion size: 501bp
- GGCAAAATAAAAAGATGTAAATTGAGACAATAAAAGCTTAAATGTAGGAGGAAAGATGGAGTTAAAGTGCAGAGTTTTTCAGTTTTTCCTTACTTGTTTGTTTCTTTTCTCTTCTTTGCAGTCAAAGTTAAGTTGTCATCAGT-L1-pA- CAAAATAACTTGTTATAACTATAAAATGTTTTGAGTAAGCT
